# Population genetic assessment of marine megafauna using seawater environmental DNA: a case study of whale sharks from the Ningaloo Coast World Heritage Area

**DOI:** 10.1101/2025.08.31.673302

**Authors:** Abinaya Meenakshisundaram, Mark Meekan, Simon Jarman, Shannon Duffy, Matilda Lamb, Laurence Dugal, W. Jason Kennington, Luke Thomas

**Affiliations:** The University of Western Australia, AIMS@UWA; The University of Western Australia; Curtin University; AIMS@UWA

## Abstract

There is a growing need for scalable and cost-effective genetic monitoring tools to support the management of species and populations of conservation importance, driven by the increasing biodiversity loss, need for long-term population data and limitations of invasive genetic sampling. Whale sharks (*Rhincodon typus*), along with other Endangered, elusive and highly migratory species, spend the majority of their lives in offshore ocean waters. This behaviour poses logistical and ethical challenges to invasive genetic sampling of the species, which is traditionally done via tissue biopsy. Here, we develop a genetic toolkit to study populations of whale sharks from seawater environmental DNA using short segments of nuclear DNA (100–300bp) containing two or more SNPs called “microhaplotypes”. Amplifying these markers from seawater collected in 1L bottles behind sharks showed that we can reliably genotype sharks from water samples taken immediately behind the animal. Moreover, we observed a strong relationship between population-level allele frequencies and estimates of genetic diversity between eDNA and tissue-derived samples, demonstrating the capability of eDNA in capturing population-level genetic information with high fidelity. We also analysed a tissue dataset of 72 sharks that attended the Ningaloo Coast World Heritage Area over six-years to showcase the utility of our SNP panel to study temporal processes in these populations. Our data revealed patterns of genetic variation through time that were consistent with whole genome techniques. Contributor estimation from mock environmental samples using various combinations of tissue-derived sequence data to estimate abundance of animals in a mixed DNA sample was shown to accurately identify the number of contributors in mixtures containing ≤ 10 individuals, beyond which biases were too large. Our findings illustrate the viability of using microhaplotype markers from seawater eDNA as a tool to study conservation genetics of whale sharks, with the potential for broader expansion of eDNA-based genetic assessments to other marine megafauna and aquatic species.

## 1. Introduction

Globally, marine megafauna are facing population declines due to various anthropogenic threats including overfishing, shipping traffic, pollution, habitat destruction, and global climate change (Costello, 2022; Germanov et al., 2018; Lewison et al., 2018; O’Hara et al., 2023; Sea et al., 2022). Population genetics has been used as a valuable tool to provide insights into the genetic diversity, population structure, connectivity, and effective population sizes of marine megafauna (Allendorf et al., 2022; Beaumont, 2007; Oleksiak & Rajora, 2019). Knowledge of such key demographic processes can help inform appropriate management measures to manage these species that play an important role in maintaining biodiversity, ecosystem health and resilience of marine environments (Carleton et al., 2013; Lewison et al., 2018; Pimiento et al., 2020).

Invasive genetic sampling of such species can be logistically complex, labour-intensive, costly, temporally limited and constrained by ethical considerations, such as minimizing harm during sampling of a protected species (Dugal et al., 2022a; Foote et al., 2012; Meekan et al., 2017; Suarez-Bregua et al., 2022). There is a critical need to develop non-invasive, affordable and robust toolkits to track individuals and populations as climate change progresses and threats to biodiversity and genetic diversity intensify (Adams et al., 2019; Foote et al., 2012).

Isolating traces of DNA left by organisms in the environment (eDNA) represents an alternative and non-invasive approach for genotyping a target species (Adams et al., 2019; Taberlet et al., 2018). Although the use of eDNA for sampling community diversity and single species detection has been adopted quickly (Adrian-Kalchhauser & Burkhardt-Holm, 2016; Berry et al., 2019; Dugal et al., 2022b; Nevers et al., 2018; Takahashi et al., 2023), the application of eDNA-based sequencing to population genetic studies remains limited but is now expanding rapidly in both terrestrial and aquatic systems (Adams et al., 2019).

Whale sharks (*Rhincodon typus*) are iconic, charismatic filter-feeding marine megafauna and the largest living fish species (up to 18 m in length(McClain et al., 2015)), central to multi-million-dollar eco-tourism industries in several tropical regions (Huveneers et al., 2017). Whale sharks have been listed as “Endangered” under the International Union for the Conservation of Nature’s Red List due to a >50% decrease in population size over the last 75 years (IUCN, 2016), likely due to anthropogenic threats including ship-strikes, overfishing and pollution (Germanov et al., 2018; Lester et al., 2020; Rowat et al., 2021; Womersley et al., 2022). These sharks are elusive and highly migratory, spending large parts of their lives in the waters of the open ocean (Arrowsmith et al., 2021; Hearn et al., 2021; Reynolds et al., 2017). Consequently, there are large gaps in our knowledge of demographics, migration routes, breeding locations and population trends, information that could be used to better inform management strategies for the species.

eDNA-based studies on whale sharks have focussed primarily on mitochondrial DNA markers from seawater collected at aggregations sites and demonstrate the potential of eDNA to obtain population-level information (Dugal et al., 2022a; Sigsgaard et al., 2016). Mitochondrial DNA can be informative and easily amplified from seawater, however shifting these efforts towards the use of nuclear markers from eDNA can provide far greater depth and accuracy of insights into trends in genetic diversity, population structure, effective population sizes and individual identification than those obtained from the analysis of mitochondrial genetic variants (Adams et al., 2019; Andres et al., 2021, 2023).

Short nuclear DNA (nuDNA) markers called “microhaplotypes” are (100 – 300 bp) fragments of DNA containing 2 or more SNPs that serve as highly informative, multiallelic markers, ideal for single sequencing detections (Kidd et al., 2013, 2014; Oldoni et al., 2019). They have been used in the field of human forensics to provide valuable data for identifying individuals in mixed DNA samples, determining biogeographic ancestry, and inferring familial relationships (P. Chen et al., 2018; Oldoni et al., 2017, 2019, 2020), and in wildlife conservation genetics to study population structures, gene flow and local adaptations through invasive genetic sampling (Baetscher et al., 2022; Hopken et al., 2023; Morin et al., 2021). Unlike microsatellites, microhaplotypes avoid issues with stutter and allelic dropout, and their multiallelic nature outperforms the limited resolution of di-allelic SNPs for mixture deconvolution and lineage analysis (Andres et al., 2023; Baetscher et al., 2018). With their compact size, high polymorphism, and suitability for degraded samples, microhaplotypes are particularly suited for eDNA-based population monitoring and wildlife conservation genetics (Adams et al., 2019; Andres et al., 2023; Morin et al., 2021).

In this study, we adopt a forensic-science based approach using short microhaplotype nuclear DNA markers and develop a toolkit to study the conservation genetics of whale sharks from seawater environmental DNA. We demonstrate the effectiveness of targeting short nuclear DNA sequence for non-invasive genotyping, population diversity analyses, and contributor estimation from complex environmental samples. Our findings demonstrate that microhaplotype markers from seawater eDNA can reliably capture individual and population-level genetic diversity and contributor estimates, offering a scalable tool for studying conservation genetics of whale sharks and other marine species.

## 2. Methods

### 2.1 Sample collection

Samples for this study were collected from Ningaloo Reef, Western Australia in May 2021 with approval from the Animal Ethics Committee of the University of Western Australia (application number: RA/3/100/1715). Individual tissue samples and corresponding seawater samples (1L Nalgene bottles) were collected from and around 33 whale sharks sampled along the 60 m depth contour (Figure 1). Snorkellers collected skin tissue biopsies from each shark using sterilized biopsy samplers mounted on a Hawaiian pole sling, and samples were immediately flash frozen in liquid nitrogen for transport and stored at –80°C prior to processing. Photographs were taken of the left and right flanks of the sharks using underwater digital or video camera for photo-identification, along with data of the shark’s length and sex, when identification was possible (Table S1). Seawater samples were collected from approximately 3 m behind each individual shark using a sterile 1L Nalgene bottle prior to tissue biopsies. Water samples were filtered using sterile single use 250 ml EZ-fit funnel filtration units with 0.22 μm filter membranes and an EZ-fit Manifold base from Merck Millipore (Merck Group, www.sigmaaldrich.com) (Kumar et al., 2020; Majaneva et al., 2018). All sharks sampled for this study were identified to be unique individuals by analysing their unique spot patterns (Meenakshisundaram et al., 2021) using the I3S Classic software (Van Tienhoven et al., 2007).

**Figure 1:**
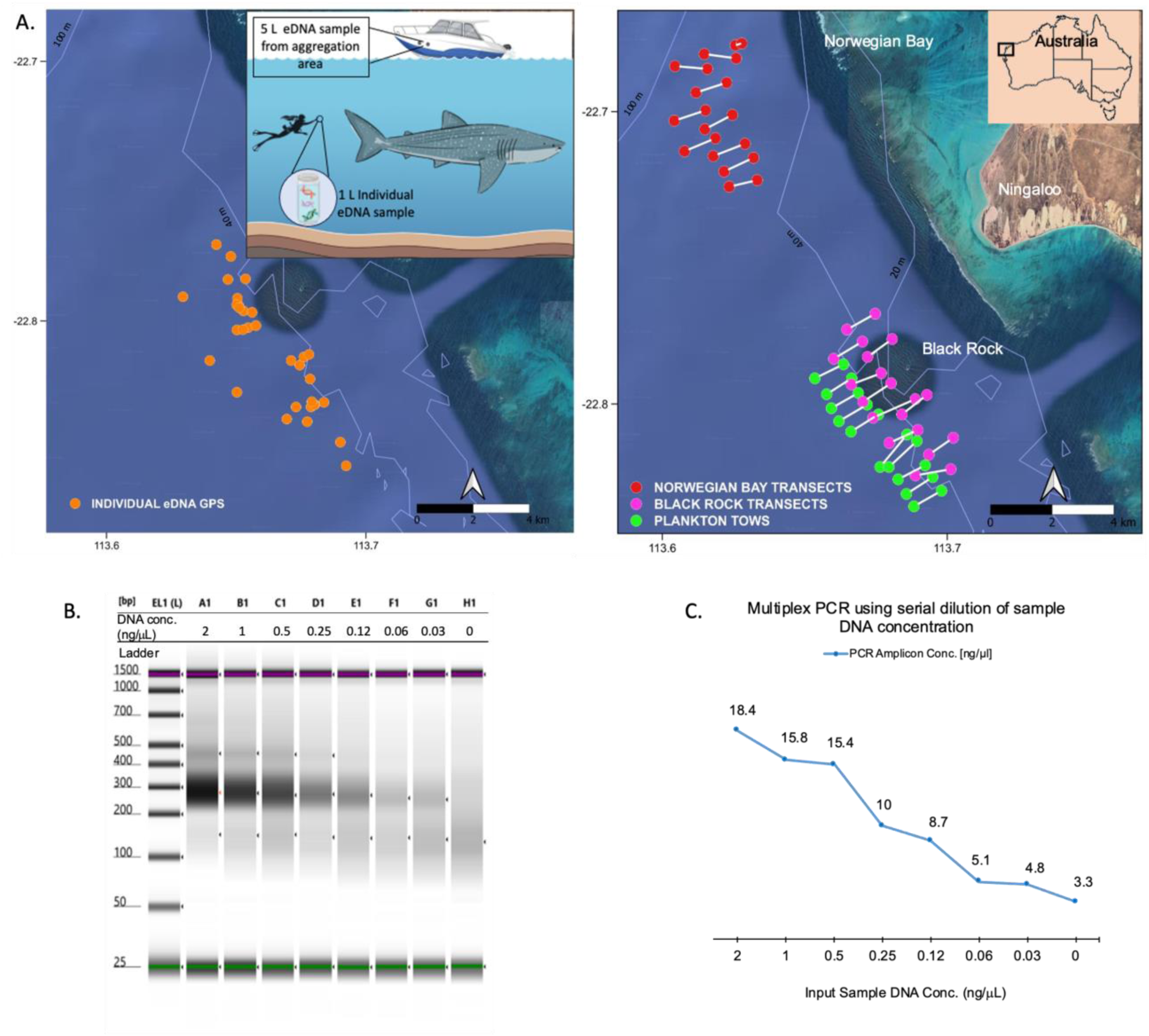
Sample collection and development of the microhaplotype panel. (A) Map of Ningaloo Reef, Western Australia showing locations of tissue and seawater collection from in-water sampling of individual whale sharks (left panel, orange points) and along 1 km transects (right panel) in the aggregation area off Norwegian Bay and Black Rock. Start and endpoints of 1 km transects where seawater samples were collected systematically every 200 m are coloured pink and red, whereas start and endpoints of transects where plankton tows operated along the entire length of the transect are coloured green. White lines show bathymetry. Outline map of Australia shows the location of Ningaloo Reef; (B) Multiplex PCR amplification of the microhaplotype panel by serial dilution of input DNA concentration visualized by TapeStation electrophoresis at target amplicon fragment sizes (∼200 – 350 bp); (C) Concentrations of Multiplex PCR products.

In addition to the water samples collected immediately behind individual sharks, seawater eDNA samples were also collected along twenty 1 km transects that were perpendicular to the reef front with locations based on GPS data from whale shark sightings during previous years (2016-2020) (Figure 1). Water was collected using a custom set-up on the sampling vessel, which included a hose fitted with weights and a mesh filter that was dropped down from the side of the boat into the ocean to a depth of 1 m and connected to a 12 V pump which directed seawater into a sterile 10 L plastic water carrier. Two litres of water were pumped into the carrier by stopping at every 200 m along the transect. The pump and hose set-up were flushed out with seawater for 5 minutes at the beginning of each transect prior to collection to prevent cross-contamination among transects. The 10 L of water collected for each transect was homogenized by shaking the container and 5 L of water was filtered using an EZ-fit Merck filtration set-up on the boat. All filter papers containing eDNA were placed in cryogenic tubes, flash frozen in liquid nitrogen for transport to the laboratory facility where they were stored at –80°C until processing.

We also sampled eDNA using towed plankton nets along 1 km transects near Black Rock (Figure 1). Tapered nets of 50 μm nylon mesh were fitted into a steel funnel that was towed alongside the boat, 0.5 – 1 m below the water surface (Alexander et al., 2023). During the tow the boat moved at two knots and sampling of the 1 km transects was usually completed in 15 minutes. The nets were then recovered and tips were cut off and stored in falcon tubes, flash frozen in liquid nitrogen for transport and kept at –80°C in the lab prior to processing.

Once filtration was completed, half of each filter paper and the tips of the tapered plankton tow nets were used for DNA extractions after cutting them into smaller pieces using sterile scissors in a laminar airflow hood that was sterilized with 10% bleach solution followed by 70% ethanol solution between each sample. DNA extraction was performed using the DNeasy Blood & Tissue kit (QIAGEN www.qiagen.com) with the addition of 360 μL ATL buffer and 40 μL proteinase K for overnight incubation at 56°C (Dugal et al., 2022a). After incubation the supernatant was extracted for DNA using a QIAcube DNA extraction robot (QIAGEN) to limit cross contamination between samples due to manual handling. Tissue samples were manually extracted for DNA using the DNeasy Blood & Tissue kit following manufacturer’s protocol. All DNA samples were stored at –80°C until use. To limit potential contamination, handling of field samples and DNA extractions took place in a separate laboratory to PCR reactions and analysis of amplicons (Goldberg et al., 2016).

A total of 50 DNA samples from whale sharks (10 samples collected each year from 2016 to 2020) used in research paper Meenakshisundaram et al. (under prep.) were used to study application of our microhaplotype marker panel to temporal patterns in population genetics.

### 2.2 Characterization of the microhaplotype panel

To identify and develop microhaplotype markers, we used DNA from 94 whale shark tissue samples for reduced representation sequencing at Diversity Arrays Technology Pty Ltd (www.diversityarrays.com). This genotyping service returned 8025 variants sites, which were then filtered to retain only sequences containing two or more SNPs. Heterozygosity was calculated for each site using the dartR package in R (Gruber et al., 2018), and only microhaplotype sequences with heterozygosity greater than 18% were retained (Kidd & Speed, 2015). The remaining target sequences were aligned to the whale shark reference genome assembly GCF_001642345.1_ASM164234v2 using the ‘blastn’ function from National Center for Biotechnology Information (NCBI) BLAST, version 2.2 (Camacho et al., 2009). Flanking regions of 100 bp on both sides of each aligned sequence were then extracted from the reference genome using BEDtools v2.31.0 (Quinlan, 2014). These extracted sequences were further aligned to the entire ‘RefSeq Genome Database’ of blastn (NCBI BLAST) to ensure species specificity, as well as to the whale shark reference genome to verify sequence specificity (Johnson et al., 2008). Primers for the selected microhaplotype sequences were designed using primer-BLAST (NCBI) (Ye et al., 2012), with melting temperatures (*Tm*) set at 55 ± 5°C, primer lengths from 18–30 bp, GC content between 40–60%, sequence specificity within the whale shark genome and resulting amplicon sizes from 162 to 251 bp. Primer pairs were checked for multiplex compatibility using PrimerPlex 2.76 (PREMIER Biosoft, http://www.premierbiosoft.com) (Kechin et al., 2020). This process led to the development of 60 primer pairs for a newly characterized microhaplotype panel, which were then ordered from Alpha DNA (http://www.alphadna.com) with added Illumina overhang adapter sequences.

### 2.3 Multiplex assay optimization

Each of the 60 primer pairs was initially tested on tissue DNA using individual PCR reactions of 10 µL that contained 5 µL of HotStarTaq Master Mix (QIAGEN), 0.2 µM of each primer (forward and reverse), and 2 ng of DNA. A two-step Touch-Down PCR (TD-PCR) protocol (Moezi et al., 2019) was applied, starting with an initial denaturation at 95°C for 15 minutes, followed by 15 cycles of denaturation at 95°C for 30 seconds, annealing at 60°C with a 0.5°C decrement per cycle for 1.5 minutes, and elongation at 72°C for 1 minute. This was followed by 40 cycles at 95°C for 30 seconds, annealing at 56°C for 1.5 minutes, and elongation at 72°C for 1 minute, with a final elongation step at 72°C for 10 minutes. The amplified products were visualized using a 4150 TapeStation system automated electrophoresis platform (Agilent technologies, www.integratedsci.com.au). Of 60 primer pairs, 40 successfully amplified the target microhaplotype amplicons at expected fragment sizes without non-target amplification.

A multiplex PCR assay was then optimized to amplify all 40 target microhaplotype markers in a single 25 µL reaction. Each reaction contained 12.5 µL of 2x QIAGEN Multiplex PCR Master Mix (QIAGEN), 0.02 µM of each of the 40 primer pairs, and either 2 ng/µL of tissue DNA or 5 µL of eDNA with DNA concentrations ranging from 0.5–3 ng/µL. The same TD-PCR conditions as the individual PCRs were used. microhaplotype markers amplified from a batch of tissue and eDNA samples using the multiplex PCR were sent for a test sequencing at Genomics WA Laboratory in Perth, Australia (www.genomicswa.com.au) using a MiSeq Nano flow cell using a 500-cycle kit (Illumina, www.illumina.com). Based on the sequencing results, 20 microhaplotype markers yielding fewer than 50 reads per marker and lacking amplification in eDNA samples were excluded from further analysis. The final optimized multiplex assay, targeting 20 microhaplotype markers expected to have 45 SNPs (Table S2), was established using the same reaction volumes and PCR conditions as previously described.

To assess the efficiency of the multiplex PCR assay for amplifying the final set of 20 microhaplotype markers from low DNA concentrations similar to those in eDNA samples, a serial dilution of tissue-derived DNA was conducted, ranging from 2 ng to 0.03 ng per PCR reaction. The PCR products from each dilution were analysed for presence of target amplicon fragments and concentration using the TapeStation electrophoresis platform and Qubit high sensitivity dsDNA kit (Thermo Fisher Scientific) respectively.

### 2.4 Library preparation

All eDNA samples were initially screened for the presence of whale shark DNA by amplifying a portion of the mitochondrial control region WSCR2-F (5′-CTATAATTGATTTAAACTGACATTTG-3′) and WSCR1-R (5′-GCATGTATAATTTTGGTTACAA-3′) primers (Castro et al., 2007) using the methodology detailed in Dugal et al. (2022a). The presence of amplification confirming presence of whale shark mitochondrial DNA at target amplicon size of ∼700 bp was verified via gel electrophoresis using the E-Gel Power Snap system (Invitrogen, Thermo Fisher Scientific). eDNA extracted from pairs of adjacent transects were pooled together and homogenized prior to library preparation.

Following amplification of 20 microhaplotype markers via multiplex PCR from both tissue and eDNA samples, samples were indexed with combinations of Illumina Nextera XT indices (Illumina) for identification. Index PCR reactions were prepared in 25 µL volumes on a QIAgility robotic workstation, minimizing the risk of cross-contamination and manual errors. Each reaction included 12.5 µL of HotStarTaq Master Mix (QIAGEN), 2.5 µL each of forward and reverse indices, and either 2 µL of tissue-derived or 5 µL of eDNA-derived multiplex PCR product to account for differences in initial DNA concentrations. The thermocycler protocol involved an initial denaturation at 95°C for 15 minutes, followed by 12 cycles of denaturation (95°C for 30 seconds), annealing (55°C for 30 seconds), and extension (72°C for 30 seconds), concluding with a final 10-minute extension at 72°C.

To generate library pools, PCR products were pooled based on equimolar concentrations that were determined using a Qubit high-sensitivity dsDNA assay. For each 96-sample PCR plate, samples were divided into two pools, dividing tissue and eDNA samples separately to equalize the concentration of DNA and tissue samples in the final pool. Primer dimers and non-target fragments were removed using a size selection step with 0.8X Ampure XP magnetic beads (Beckman Coulter, www.beckmancoulter.com). These cleaned pools were then combined at equimolar concentrations to form the final library pool, which was verified for the presence of target fragment sizes using the 4150 TapeStation system (Figure S1). The finalized library was submitted to the Genomics WA Laboratory and sequenced using an Illumina MiSeq Nano Kit (500 cycle).

### 2.5 Data analysis

#### 2.5.1 Sequence data processing

The demultiplexed sequence data, obtained as fastq files (File S1), underwent initial quality control processing with fastp-0.23.4, which removed low-quality bases, trimmed adapters, and removed poly-G tails (Chen et al., 2018). For paired-end read merging, VSEARCH was employed with a threshold allowing up to five mismatches within the overlapping regions (Rognes et al., 2016). Each of the 20 expected microhaplotype amplicon sequences was used as a reference for mapping sequenced reads (Table S2). These reference amplicons were compiled in a fasta file and indexed using samtools-1.12 (Li et al., 2009). Merged reads were mapped to these references using BWA-MEM (Li, 2013), generating BAM files that were filtered for mapping quality above 10, then sorted and indexed with samtools-1.12 (Li et al., 2009). SNP calling from the resulting BAM files was performed using bcftools-1.3.1 (Danecek et al., 2021), utilizing ‘mpileup’ and ‘call’ functions to exclude indels and call SNPs with an average depth >10 across samples. Further filtering was done with vcftools 0.1.16 (Danecek et al., 2011) to retain SNPs in Hardy-Weinberg Equilibrium (HWE) with a p-value ≥ 0.05. Ultimately, this process identified 56 SNP sites across 20 microhaplotypes (Table S2), shared between tissue and eDNA derived sequences.

#### 2.5.2 Comparison of tissue versus eDNA derived SNP data

SNP allele frequency data obtained from eDNA in seawater collected directly behind individual whale sharks were compared with data from corresponding tissue samples of each individual. To minimize sequencing and PCR artifacts, alleles with fewer than five reads were excluded from all samples. Read counts were then normalized to 100 reads per sample, accounting for variations in initial DNA concentrations and sequencing depths. Additionally, alleles with fewer than 0.01 scaled reads were filtered out of the dataset.

Population allele frequency estimates derived from tissue and eDNA samples were used to assess the accuracy of population genetic diversity estimates obtained from eDNA. Significance of the correlation between population allele frequencies from eDNA and tissue was evaluated using Spearman’s correlation statistic (ρ) with the ‘cor.test’ function from R package stats (Ali Abd AlHameed, 2022; R Core Team, 2022) for the most common allele at each locus.

SNP allele frequencies were calculated for each individual sample to evaluate the concordance between genetic profiles derived from eDNA and those obtained from corresponding tissue samples. This comparison aimed to determine the reliability of eDNA in accurately reflecting individual genetic profiles. Alleles with frequencies less than 1% were removed from the data set to avoid sequencing errors. Principal components analysis (PCA) was performed with the ‘prcomp’ function of R package stats (R Core Team, 2022). A dendrogram based on Euclidean genetic distances between samples was generated using ‘dist’ and hierarchically clustered using the R package stats-4.3.1 (R Core Team, 2022).

#### 2.5.3 Temporal population genetics using microhaplotype markers

To explore the utility of this panel to study the temporal genetics of whale shark aggregations, we used microhaplotype markers to analyse tissue samples from 50 individual whale sharks spanning 2016 to 2020, along with 22 paired tissue and eDNA samples collected in 2021. Estimates of observed heterozygosity (Ho), unbiased expected heterozygosity (uHe), Inbreeding coefficient (*F*_IS_) were calculated with the function ‘gl.report.heterozygosity’ of R package dartR (Gruber et al., 2018) and allelic richness (AR) were calculated using ‘allel.rich’ function of package PopGenReport (Adamack & Gruber, 2014). These estimates were compared to those obtained from whole-genome genetic variants presented in the study Meenakshisundaram et al. (under prep.). A paired Wilcoxon signed rank exact test implement using the ‘wilcox.test’ function of R (R Core Team, 2022) was used to test for differences in uHe and AR estimates obtained using whole-genome genetic variants versus microhaplotypes. Additionally, genetic diversity estimates from eDNA samples were compared to those obtained from corresponding tissue samples to examine the efficacy of eDNA-based genetic monitoring.

Principal component analysis (PCA) was performed using the ‘snpgdsPCA’ function of the R package SNPRelate-1.34.1 (Zheng et al., 2012).

#### 2.5.4 Simulated contributor estimation

Probability of identity (*P_ID_*) as well as probability of identity between siblings (*PI_sibs_*) were calculated using function ‘pid_calc’ from R package PopGenUtils to evaluate the efficiency of this set of microhaplotype markers for individual identification and mixture deconvolution (Taberlet & Luikart, 1999).

Simulated complex DNA samples were made by pooling allele frequency estimates from individual tissue samples in combinations of 1, 3, 5, 8, 10, 20, 30, 40, 50, and 80 individuals. All alleles with less than 0.01 frequency in individual samples were removed to reduce sequencing errors (Andres et al., 2021; Meenakshisundaram et al., 2025). Similarly, simulated DNA mixtures were made by pooling allele frequencies estimated from eDNA samples collected behind individual whale sharks in combinations of 1, 3, 5, 8, 10 and 20.

To estimate the number of genetic contributors in a mixed sample, we employed a likelihood-based mixture model (Egeland et al., 2003; Sethi et al., 2019). This approach provides likelihood estimates of the number of contributors (x) based on the observed alleles (A = {*a_1_,…,a_n_*}) and their corresponding population allele frequencies (*p =* {*p_1_,…,p_n_*}) for each locus (*j*) using:

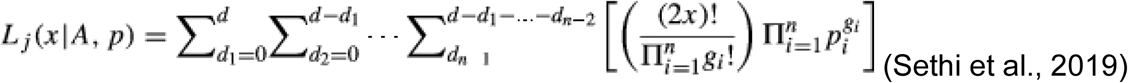

The maximum likelihood estimate of the number of contributors is determined by the product of these likelihoods across all loci. A custom R-script made by Andres et al. (2021) was modified for mixture deconvolution using SNP data (Meenakshisundaram et al., 2025). We varied the number of individuals included in the analysis to evaluate the potential benefits of utilizing larger sample sizes for estimating total population allele frequencies used for estimating abundance (Andres et al., 2021). The putative number of individuals (*y*) provided to the model corresponded to the maximum number of individuals present in any mixture within the dataset: 80 for tissue simulated mixtures and 20 for eDNA simulated mixtures.

## 3. Results

### 3.1 Results from microhaplotype amplification and sequence data

Multiplex PCR amplification of the panel of 20 microhaplotype markers by serial dilution of input of tissue-derived DNA concentrations showed a gradual decrease in concentration and visualization of PCR products, with minimal levels of amplification occurring at low DNA concentrations of 0.03 to 0.06 ng/μL (Figure 1). This demonstrates the potential applicability of this multiplex assay for analysis of eDNA samples from seawater, which generally contain low concentrations of target DNA.

Screening of eDNA samples collected behind individual whale sharks for the presence of whale shark mitochondrial DNA (mtDNA) by amplifying a portion of the WSCR region resulted in successful amplification in 25 out of 33 samples (Figure S2). Among the samples that did not yield amplification for this mtDNA region (sample IDs 15, 31, 35, 45, 51, 53, 56, and 64), three samples (15, 31, and 45) did amplify nuclear microhaplotype markers, with an average read count of 572 (± 455 SD). Conversely, five samples that showed successful mtDNA amplification lacked sufficient amplification of nuclear microhaplotype markers. eDNA samples collected from transects across the aggregation area showed no evidence of whale shark mtDNA, except for low levels of amplification in samples 5 and 6 from Black Rock transects (Figure S2). No amplification or sequence data for the microhaplotype panel was obtained from any transect sample.

High average read depth per sample was observed across 33 tissue samples (9370.8 (± 802.5 SD)) collected in 2021, whereas, for the corresponding eDNA samples, the average read counts per sample was substantially lower (260.4 (± 223.2 SD)) (Figure S3). The 56 SNPs called from tissue samples were found to have mean depth per site of 168.6 (± 800.5 SD) (Figure S3), average per-SNP quality of 2299.2 (± 2946.8 SD). Of the 33 eDNA samples, 22 retained sufficient data over five or more SNP loci after filtration and were included in further analyses alongside their corresponding tissue samples. The same variant positions called from the selected 22 eDNA samples had a mean depth of 5.9 (± 205.2 SD) per site (Figure S3), average per-SNP quality of 2299.2 (± 2946.8 SD).

Genetic variants were called from 50 temporal whale shark samples collected through years 2016 to 2020 using positional data obtained from the 56 SNPs found common to both 2021 tissue and eDNA sequence data. This retrieved 48 of the 56 SNPs in this dataset. All negative controls included during DNA extraction and PCR steps did not produce amplification, which was checked using a TapeStation electrophoresis platform, thus confirming absence of cross-contamination.

### 3.2 Genotyping sharks from water samples

We were able to reliably genotype individual sharks from seawater samples using a panel of microhaplotype markers targeting nuclear DNA. Allele frequency data was successfully obtained for 58 out of 112 alleles across both eDNA and tissue-derived samples. A strong and statistically significant correlation was observed between population allele frequencies derived from eDNA and tissue samples, with a Spearman’s correlation coefficient (ρ) of 0.81 and a p-value of 1.2e-14 (Figure 2A). Allele frequencies were obtained for a range of 6 to 39 loci (average = 18.8, SD = 9.6 loci) across each water sample, with corresponding loci retained from the matching tissue sample data. The PCA plot (Figure 2B) revealed that eDNA and tissue samples from the same individual clustered closely in multidimensional space, indicating high similarity in allele frequencies. This clustering pattern was further confirmed by a dendrogram constructed using Euclidean genetic distances, where samples from each individual whale shark consistently showed least distances and clustered tightly (Figure 2C). However, when the analyses were repeated by retaining allele frequencies of all 112 alleles from tissue samples, the patterns of grouping in PCA and Euclidean distance plots was based on sample type (eDNA or tissue) rather than by individual (Figure S4).

**Figure 2:**
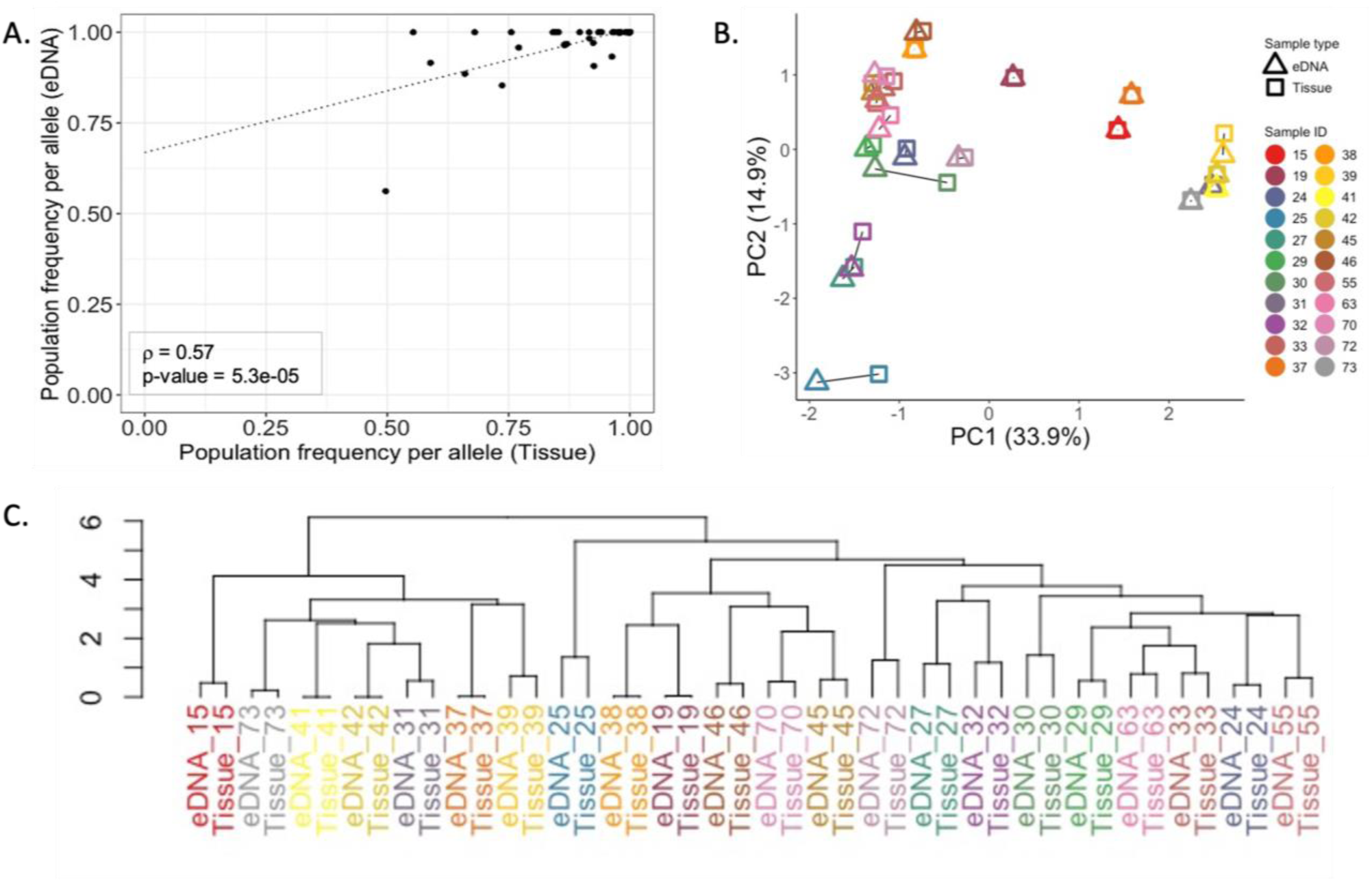
(A) Scatter plot showing the correlation between population allele frequencies for SNP alleles obtained from eDNA and tissue-derived sequences, with statistical significance indicated by Spearman’s correlation coefficient (ρ); (B) Principal Component Analysis (PCA) plot of individual eDNA and tissue samples, colour-coded by sample ID, with solid black lines connecting sample pairs that correspond to the same individual shark, for visualisation of individual-level clustering; (C) Dendrogram illustrating the hierarchical clustering of samples based on Euclidean distances colour-coded by sample ID.

### 3.3 Population genetic inferences using microhaplotype markers

For sharks sampled between the years 2016 to 2021, the microhaplotype panel (48 SNP sites) revealed consistent Ho values ranging from 0.151 to 0.231, with an average of 0.192 (± 0.03 SD) (Table S3) and similarly varying uHe values with an overall mean of 0.173 (± 0.02 SD) (Figure 3; Table S3). Consistent AR values averaging 1.17 (± 0.02 SD) (Figure 3; Table S3) and negative Inbreeding coefficients (Table S3) suggested a healthy level of genetic diversity and minimal inbreeding within the population over the study period. In comparison, genetic diversity estimates derived from lcWGS (7585 SNP sites) (Meenakshisundaram et al.,under prep.) were higher, with average Ho value of 0.22 (± 0.007 SD) (Table S3), uHe 0.192 (± 0.007 SD) and AR 1.82 (± 0.03 SD) (Figure 3; Table S3). Estimates of uHe obtained from both methods showed no significant differences (p-value = 0.06); whereas, AR estimates obtained from microhaplotypes were significantly lower than those obtained from lcWGS (p-value = 0.03). uHe values were slightly lower than Ho values and inbreeding coefficients were negative, similar to those obtained from microhaplotype markers, suggesting similar population dynamics. In 2021, genetic diversity metrics estimated using eDNA samples collected from the same individuals as the tissue samples aligned closely with those obtained from tissue samples, with slightly lower uHe and AR values in eDNA-derived estimates (Figure 3). The PCA plot (Figure 3) did not show observable variation in microhaplotype marker derived SNP data between samples collected during different years, with a few outliers observed in the eDNA cohort.

**Figure 3.**
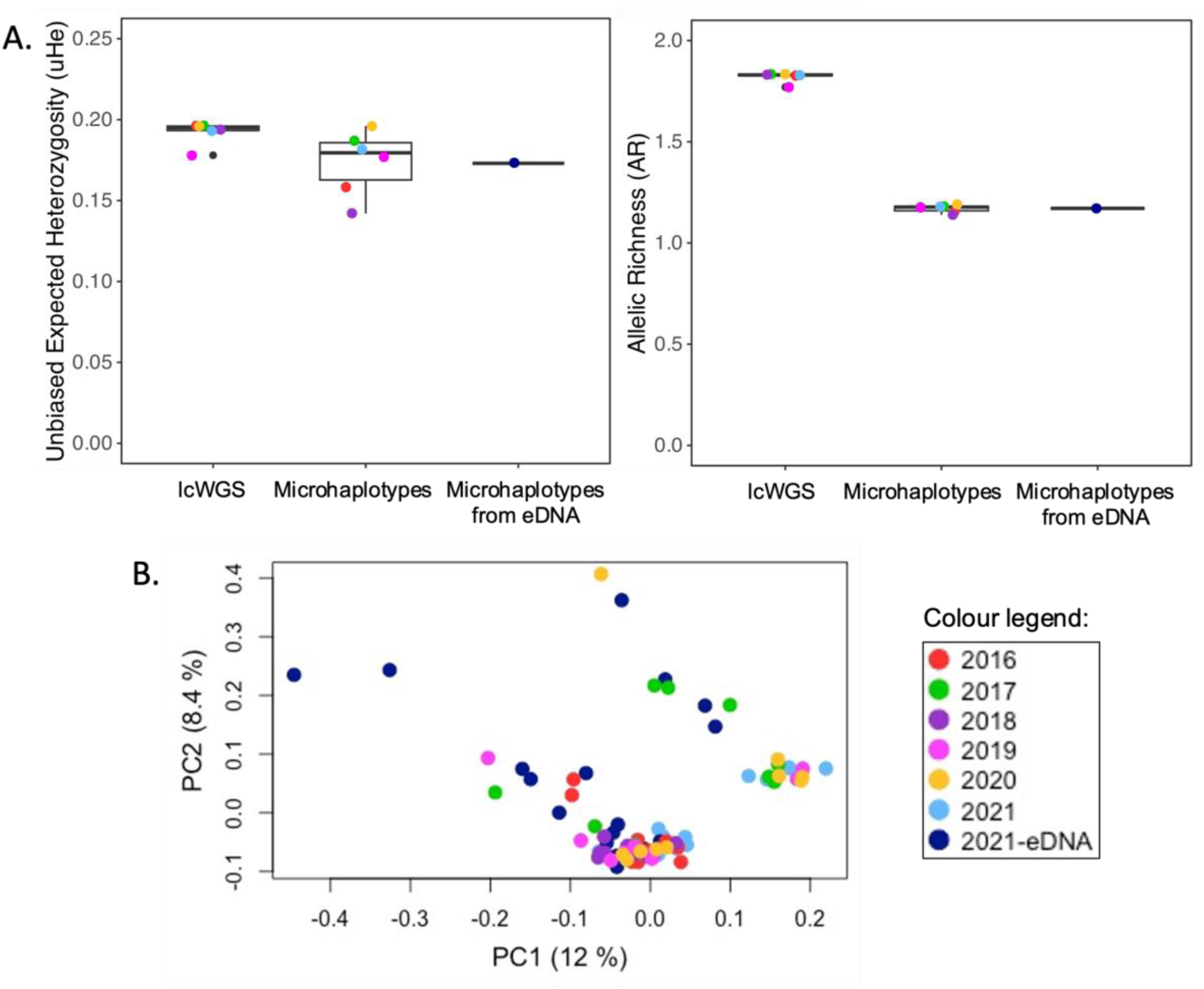
(A) Unbiased expected heterozygosity (uHe) and allelic richness (AR) of whale sharks attending the Ningaloo Reef aggregation from 2016 to 2021, estimated from tissue samples using genome-wide SNPs derived from low-coverage whole-genome sequencing (lcWGS) and microhaplotypes. Genetic diversity estimates from microhaplotypes obtained from environmental DNA (eDNA) samples collected in 2021 are also included for comparison; (B) Principal component analysis (PCA) plot illustrating genetic variation among microhaplotypes derived from whale shark tissue samples (2016–2021) and eDNA samples collected in 2021. Data points are color-coded by year and sample type.

### 3.4 Simulated contributor estimation from mixed samples

The combined probability of identity (*P_ID_*) for all 56 loci was estimated at 6.04 × 10⁻⁹ using tissue-derived SNP data and 1.93 × 10⁻⁸ using individual eDNA-derived SNP data, with *PI_sibs_* values of 7.05 × 10⁻⁵ and 1.46 × 10⁻⁴, respectively, demonstrating the effectiveness of this microhaplotype panel for individual identification and mixture deconvolution. For DNA mixtures simulated using tissue-derived sequence data, contributor estimations derived using population allele frequencies from 83 individuals closely matched true contributor numbers when the mixture contained up to 10 individuals. However, as the true contributor count increased, so did the discrepancies, and mixtures with higher true contributor counts of 20, 30, 40, and 50 exhibited substantial underestimations. A similar pattern emerged when using population allele frequencies from a smaller sample of 33 individuals (Figure 4). At lower true contributor counts, the model showed higher accuracy, with minimal or no bias in estimating 1, 3, 5, and 10 contributors, yielding biases of 0, –1, +1, and 0, respectively. However, at higher true contributor counts, negative biases increased: for 20, 30, 40, 50, and 80 contributors, estimates deviated by –28, –25, –16, –45, and – 18, respectively. These findings suggest that although the maximum likelihood method reliably estimates contributor numbers in mixtures with low true counts, bias increases with higher true contributor counts, particularly when population allele frequencies are based on smaller sample sizes (Figure 4).

**Figure 4.**
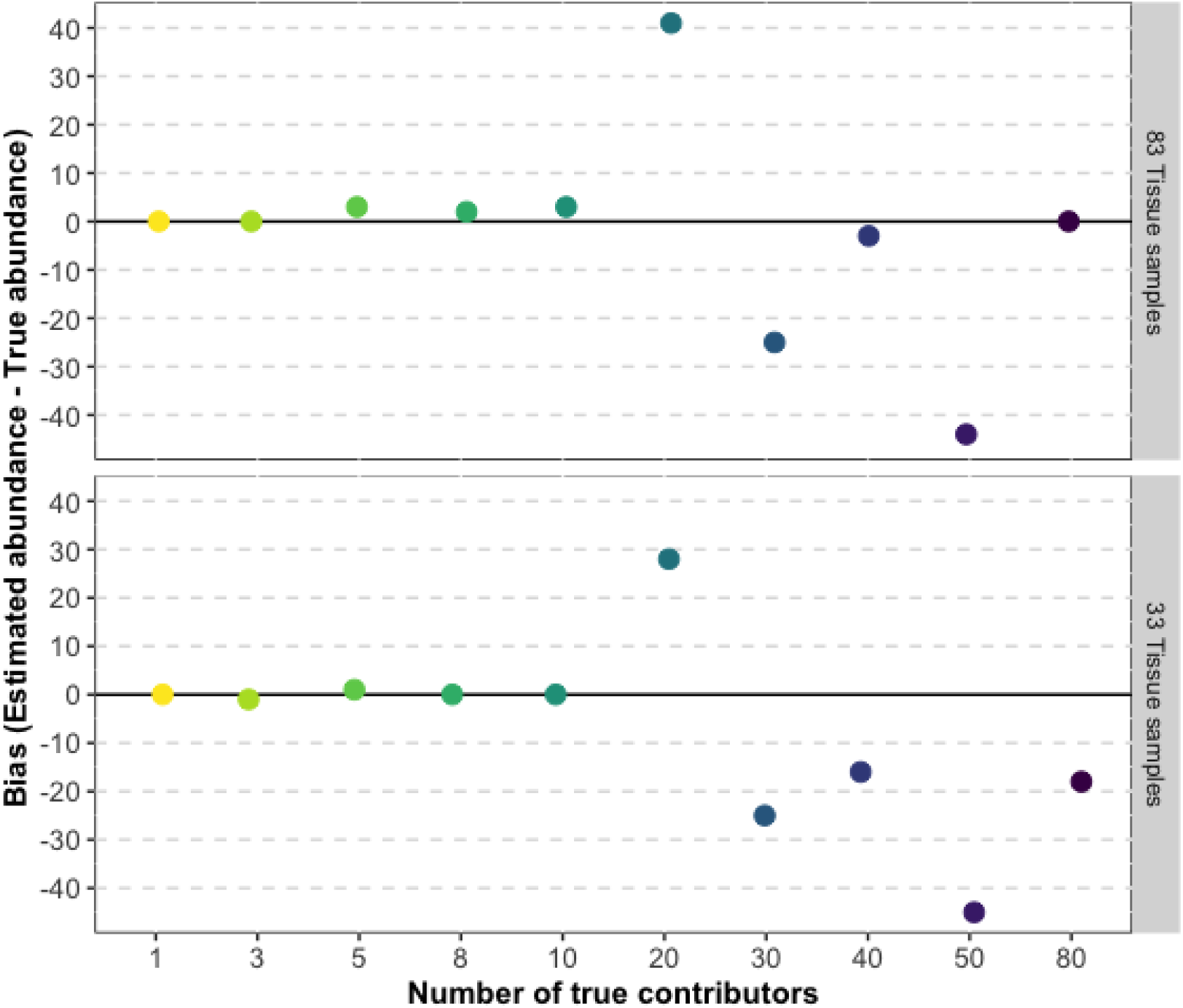
Bias in number of contributors estimated using maximum likelihood approach from simulated whale shark DNA mixtures made using individual tissue-derived sequence data, colour-coded by the known number of individuals. The abundance estimation was performed using population allele frequencies calculated using tissue-derived sequence data from varying number of individuals as denoted on the right of each grid.

Estimation of the number of contributors in simulated DNA mixtures made using eDNA-derived sequence data from samples collected near individual whale sharks was analysed using population allele frequencies derived from both tissue and eDNA sources (Figure 5). All estimation models produced accurate results when the true number of contributors was 1, but accuracy declined for higher contributor numbers when compared to estimates obtained from tissue-derived simulated mixtures.

**Figure 5.**
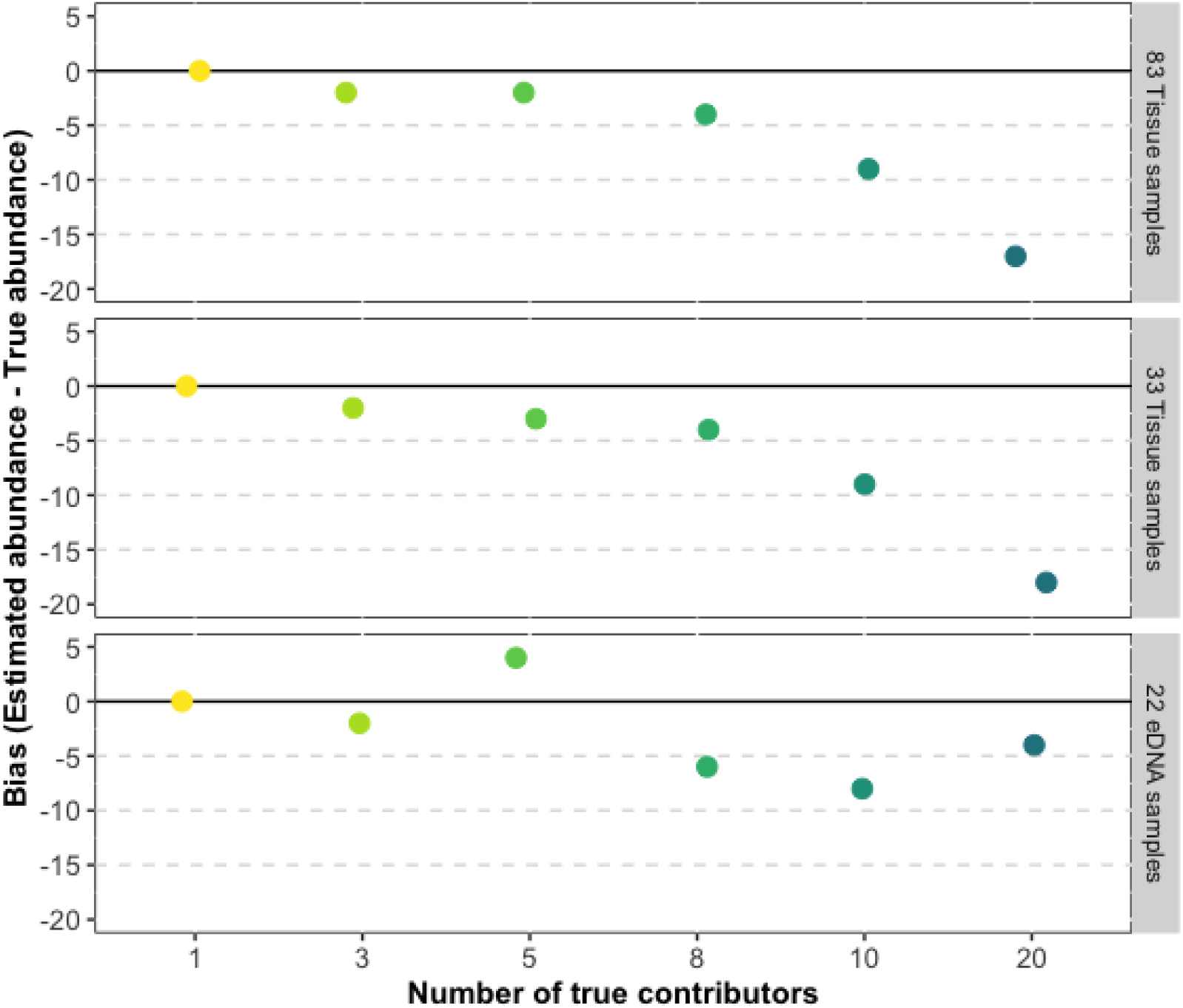
Bias in number of contributors estimated using a maximum-likelihood model from simulated DNA mixtures of whale sharks made using individual eDNA-derived sequence data, colour-coded by the known numbers of individuals. The abundance estimation was analysed using population allele frequencies calculated using tissue or eDNA-derived sequence data from varying numbers of individuals as shown on the right of each grid.

Using population allele frequencies from 83 tissue samples, the model showed slight negative biases of –2 for true contributor counts of 3 and 5, with bias increasing as the true contributor count rose. A similar trend was observed when using allele frequencies from 33 tissue samples, although the negative bias was marginally higher, increasing by approximately –1 individual compared to estimates based on the larger 83-sample dataset.

When population allele frequencies were derived from 22 eDNA samples, the model showed no bias when the true count was 1 (bias = 0) and a bias of –2 when the true count was 3. For higher contributor counts of 10 and 20 individuals, a lower negative bias in estimates was observed than those obtained using tissue-derived population allele frequencies.

## 4. Discussion

Our results demonstrate that multi-allelic nuclear DNA (nuDNA) markers sequenced from seawater environmental DNA can effectively be used to genotype individuals and capture population-level allele frequency data of populations. Individual-level clustering patterns based on genetic distances between eDNA and tissue samples confirm that eDNA derived from seawater collected in close proximity to whale sharks can accurately reflect the genetic signatures of specific individuals. Additionally, metrics of genetic diversity from eDNA samples were closely aligned with tissue-derived estimates, with only slight reductions in allelic richness and heterozygosity. The maximum-likelihood approach for estimating contributor numbers in complex multi-genotype DNA mixtures yielded accurate estimates for low contributor counts (<10 individuals), though biases increased as the complexity of the sample increased. These findings underscore the utility of eDNA-based techniques for reliable genotyping, diversity assessment, and abundance estimation for marine megafauna, and provides a scalable, ethical alternative to invasive genetic sampling. We had variable success in amplifying whale shark mtDNA and nuclear microhaplotype markers from seawater samples collected near individual whale sharks. Although mtDNA failed to amplify in five samples that successfully yielded nuclear microhaplotype markers, three samples amplified mtDNA but not nuDNA, suggesting differential presence and/or degradation of DNA due to environmental, molecular, species or sampling factors. Although mtDNA is often preferred in marine studies due to its high copy number per cell, which typically increases initial abundance and detectability (Dejean et al., 2011; Marshall & Parson, 2023), it can degrade at rates comparable to or even faster than nuDNA under certain environmental conditions, such as increased microbial activity and oxidative stress (Bylemans et al., 2018; Jo et al., 2022; Moushomi et al., 2019; Shokolenko et al., 2009). However, our findings suggest that both types of DNA have variable detection, emphasizing the need to consider sample-specific degradation (Hughes-Stamm et al., 2011) and possibly the inclusion of both mtDNA and nuDNA markers in studies of marine eDNA.

A significant discrepancy in read depth was observed between tissue and eDNA samples, with the latter yielding substantially lower average read counts of 260.4 (± 223.2 SD) per sample. Although sufficient for identifying individual genetic signatures and estimating population-level allele frequencies, the read depth of 6 to 39 SNPs recovered from eDNA samples fell short of the 56 SNPs identified in tissue samples, resulting in lower estimates of genetic diversity and insufficient data for mixture deconvolution tasks. In comparison, Andres et al. (2021) reported an average read depth of 4,305 reads per sample for nuclear DNA microsatellite markers sequenced from water samples collected in areas of high densities of round gobies (*Neogobius melanostomus*), highlighting the challenges of achieving adequate read depth with nuclear markers in eDNA studies.

Notably, much higher read depths have been achieved in eDNA studies that targeted mitochondrial genomes. For example, Sigsgaard et al. (2016) and Parsons et al. (2018) reported average read depths of 263,111 and 237,434.5 reads per sample for seawater samples collected near whale sharks and from harbor porpoise (*Phocoena phocoena*) fluke prints, respectively. These differences likely stem from sequencing platform choices and associated read capacity; our study used the MiSeq Nano kit with a capacity of around 1 million reads per run, which is substantially lower than the MiSeq v2 kit (up to 15 million reads per run) (Andres et al., 2021), Miseq v3 kit (up to 25 million reads per run) (Sigsgaard et al., 2016) and the HiSeq platforms (up to 4-6 billion reads per run) (Parsons et al., 2018) used in such studies. For future work, the use of higher-capacity sequencing kits would improve read depth per sample, enhancing the resolution of eDNA for population-level analyses and enabling more robust applications in genetic diversity assessment and mixture deconvolution. Additionally, employing technical replicates during PCR (Shirazi et al., 2021) and implementing unique molecular identifiers (Staadig et al., 2023) to minimize PCR biases and errors to improve data quality and accuracy can be incorporated in future studies.

Beyond improvements in molecular techniques, increasing the water sample volume, improving filtration methodology and limiting degradation during sample transport can also improve eDNA yield. For open marine environments where DNA is more diluted, increasing sample volumes from the typical 1-2 L to 4-5 L has been shown to enhance DNA capture (Fraija-Fernández et al., 2020). In this study, 1 L seawater samples were collected in proximity to each whale shark, held in enclosed ice boxes for 3 to 8 hours prior to filtration and storage at –80°C. Implementing field-based filtration systems with larger filters, such as enclosed Sterivex filter cartridges (Merck Group), could further enhance results by enabling the filtration of larger water volumes (∼4 L). These enclosed filters not only retain more eDNA but also protect the filtered DNA from contamination, preserving its integrity during transport (Miya et al., 2016; Wong et al., 2020).

eDNA samples and tissue samples from the same individuals exhibited high correlation in allele frequencies, suggesting the possibility of conducting large-scale population genetic assessments based on eDNA alone, allowing for the identification and tracking of individual whale sharks without the need for invasive tissue collection.

This is consistent with observations made in the study by Dugal et al. (2022a), which showed that mitochondrial eDNA haplotyping from seawater eDNA collected behind whale sharks matched tissue-derived haplotypes with 100% sequence accuracy. Obtaining accurate allele frequencies from individual organisms using seawater eDNA enables further genetic analyses such as intraspecific genetic variation, population structure, genetic differentiation. In addition, analyses such as kinship, relatedness, inbreeding and population assignment tests can be done on par with tissue-derived sequence data, avoiding the need for specialized bioinformatic approaches required for samples containing DNA from multiple individuals (Adams et al., 2019; Sigsgaard et al., 2020a). However, this targeted approach is best suited to sparse, low-density whale shark aggregations such as Ningaloo Reef and Maldives (M. Meekan et al., 2006; Riley et al., 2010). Localized and high-density gatherings, such as those off the Yucatan coast of Mexico (De La Parra Venegas et al., 2011) and Qatar (Sigsgaard et al., 2016), might yield overlapping signals from multiple individuals. Additionally, the persistence of eDNA in the marine environment, which can extend up to several days (Collins et al., 2018; Thomsen et al., 2012), could make the unambiguous detection of individual genetic signatures more challenging. To address these challenges, studies should consider population densities and the biological traits of target species (e.g., social structure, breeding behaviours) that may influence eDNA distribution.

These outcomes emphasize the accuracy of multi-allelic nuclear markers derived from eDNA for non-invasive genetic analyses of populations, even from complex samples. Although it is challenging to assign individual genotypes from DNA samples that contain a mixture of individuals, bioinformatic frameworks designed for pooled sequencing data (Pool-seq) can enable robust genetic analysis of populations (Gautier et al., 2013). Tools that adjust for skewed allele frequencies in DNA samples from a mixture of individuals that arise from unequal individual contributions, variable sequencing depth, and loss of rare alleles, allow estimation of key parameters such as genetic diversity, inbreeding, population structure, and kinship (Czech et al., 2024; Gautier et al., 2013, 2022; Sigsgaard et al., 2020a). Although the technique used for eDNA collection in transects across Ningaloo Reef did not show amplification of microhaplotype markers, this methodology could prove useful for studies of genetics in aggregations where there are localized high-densities of whale sharks.

Estimates of genetic diversity derived by amplifying microhaplotype markers from tissue samples collected from years 2016 to 2021 aligned closely with the results obtained from the lcWGS data, indicating that the microhaplotype panel provided a reliable measure of genetic variation in the population of whale sharks at Ningaloo. Both marker sets showed little temporal variation in allelic richness, stable expected heterozygosity values and negative inbreeding coefficients, indicating the presence of a population with stable genetic diversity and no signs of inbreeding. The higher observed than expected heterozygosity and lack of significant inbreeding observed in both studies could also suggest genetic admixture (Boca et al., 2020), owing to the maintenance of stable genetic diversity in the population, despite population declines observed globally. Similar observations of stable patterns in genetic diversity have also been observed in other aggregations of whale sharks across the Indo-Pacific (Castro et al., 2007; Hardenstine et al., 2022; Schmidt et al., 2009). Such results and our results here contradict the hypothesis that there was a decline in the genetic diversity of the population of whale sharks in Western Australia from 2007 to 2012 (Vignaud et al., 2014).

The whale sharks found at Ningaloo Reef are a part of a seasonal feeding aggregation that is dominated by juvenile males (males to females abundance ratio = 4:1) (Meekan et al., 2006; Meekan et al., 2020). Globally, most aggregations display similar biases in the sex and size of animals (Rohner et al., 2021), with the exception of a few that are dominated by females (Acuña-Marrero et al., 2014; Ketchum et al., 2013). It is thus important to consider that the individuals sampled from aggregations may not fully represent broader populations.

Despite concerns about the impacts of anthropogenic pressures (IUCN, 2016), to date no genetic signature of population declines have been detected. The potential longevity (up to 100+ years) (Bradshaw et al., 2007; Meekan et al., 2020) and long generation times of whale sharks may buffer against rapid declines in genetic diversity (Lippé et al., 2006), making it challenging to identify the effects of recent population bottlenecks. This highlights the need for ongoing, long-term monitoring to assess genetic trends in this Endangered species. Non-invasive techniques using seawater eDNA methodology outlined here could complement traditional methods for stable long-term monitoring of, supporting informed management decisions (Yoccoz, 2012).

Estimations of individual contributors from simulated DNA mixtures based on tissue-derived sequence data demonstrated high accuracy for mixtures containing low numbers of individuals (1, 3, 5, 8, 10), with increasing negative bias as number of contributors rose. However, simulated mixtures using eDNA-based data lacked sufficient read depth and SNP coverage to support reliable estimation of abundance. This result contrasts with an earlier eDNA study of zebrafish conducted in controlled aquaria that accurately estimated mixtures of up to 20–25 individuals using a 69-SNP microhaplotype panel (Meenakshisundaram et al., 2025). Additionally, a simulation study by Andres et al. (2023) using genetic data from linesnout goby (*Elacatinus lori*) (D’Aloia et al., 2020) found that a 64-SNP panel effectively resolved mixtures up to 25 individuals. However, eDNA samples from ocean environments often suffer from lower read depth and loss of rare alleles due to factors such as UV exposure, salinity, microbial degradation, temperature fluctuations, and variable eDNA dispersion. These can contribute to underestimations and false negatives in abundance estimates (Collins et al., 2018; Murakami et al., 2019; Tsuji et al., 2017). Using total population allele frequencies derived from a larger sample set in the calculation for estimating abundance reduced negative bias for higher contributor counts (>10 individuals) and similar results have been observed in previous studies using this maximum-likelihood framework (Andres et al., 2023). Nevertheless, this did not significantly improve estimates for smaller contributor numbers, indicating that abundance estimation for marine megafauna typically found in smaller numbers can be effectively performed using a database of sequence data from limited number of individuals.

In conclusion, this study demonstrates that microhaplotype markers sequenced from seawater eDNA provide a reliable, non-invasive alternative for genetic monitoring of whale sharks, at scales of both individuals and populations. The methodology we developed offers a valuable tool for assessing intraspecific genetic variation and abundance in marine megafauna and other aquatic species that are typically sparsely distributed, such as cetaceans, sharks, turtles and dugongs (James et al., 2005; Rosenbaum et al., 2009; Rus Hoelzel et al., 2006; Sigsgaard et al., 2020b). This approach is especially advantageous for monitoring highly migratory species that occupy expansive habitats or are otherwise challenging to sample invasively due to logistical or ethical considerations (Dekker, 2016; Gero et al., 2014; Welsh et al., 2008). With further optimization, this eDNA-based technique could potentially be applied to a broader range of species, reducing the reliance on physical captures and invasive sampling for future population genetic studies. Overall, our study offers a promising framework for expanding eDNA applications in genetic monitoring, supporting conservation efforts for whale sharks and other vulnerable marine megafauna species across diverse global ecosystems.

## Acknowledgements

We acknowledge the West Thalanyji people, the traditional custodians of the land and water at Nyinggulu on which field work was conducted, and pay our respects to their Elders past, present and emerging. This project was supported and funded by the Minderoo Foundation through the Minderoo Foundation Exmouth Research Laboratory (MERL) and its staff. Tissue sampling and field data collection for this study was conducted as a part of an annual long-term monitoring study on the Ningaloo whale sharks by Australian Institute of Marine Science (AIMS) supported by Santos Limited and we gratefully acknowledge all members of the research team involved in this field work. We also thank the Genomics WA Laboratory in Perth, Australia for sequencing.

## Author contributions

Authors AM, SJ and LT designed the study. AM and LT obtained funding for the project. AM, SD, LT and MM undertook sample collection. AM undertook laboratory work, data analysis and drafting of manuscript. ML assisted with laboratory work and LD provided WSCR primers and guidance for associated lab protocols. All authors contributed to writing of the manuscript.

## 4.10 Supplementary Information

**File S1.** Link to raw fastq sequence data for all samples used in the study, DArT SNP data, reference microhaplotype amplicon sequences and the script used for SNP calling and bioinformatic analysis ()

**Table S1.** Metadata of the individual whale sharks that were sampled for both eDNA and tissue biopsies.

| Sample ID | Date of collection | Length of shark (m) | Sex | Latitude | Longitude |
| --- | --- | --- | --- | --- | --- |
| 15 | 08-May-21 | 5.5 | M | -22.7990 | 113.6522 |
| 16 | 08-May-21 | 5 | F | -22.7998 | 113.6509 |
| 17 | 08-May-21 | 8 | M | -22.7991 | 113.6520 |
| 18 | 08-May-21 | 6 | M | -22.7980 | 113.6512 |
| 19 | 08-May-21 | 6 | M | -22.7982 | 113.6554 |
| 22 | 08-May-21 | 6.5 | F | -22.7901 | 113.6506 |
| 24 | 08-May-21 | 7 | M | -22.7853 | 113.6470 |
| 25 | 08-May-21 | 4.5 | F | -22.7825 | 113.6484 |
| 27 | 08-May-21 | 3.5 | M | -22.7816 | 113.6502 |
| 28 | 09-May-21 | 5 | M | -22.7837 | 113.6440 |
| 29 | 09-May-21 | 4 | F | -22.7947 | 113.6252 |
| 30 | 09-May-21 | 5 | M | -22.7837 | 113.6158 |
| 31 | 09-May-21 | 4.5 | M | -22.7731 | 113.6252 |
| 32 | 09-May-21 | 7.5 | M | -22.7723 | 113.6291 |
| 33 | 09-May-21 | 5 | M | -22.7730 | 113.6273 |
| 35 | 09-May-21 | 3.5 | F | -22.7665 | 113.6275 |
| 37 | 09-May-21 | 5 | F | -22.7645 | 113.6258 |
| 38 | 09-May-21 | 7 | M | -22.7621 | 113.6254 |
| 39 | 09-May-21 | 6 | F | -22.7554 | 113.6282 |
| 40 | 09-May-21 | 8 | M | -22.7476 | 113.6231 |
| 41 | 09-May-21 | 7 | M | -22.7435 | 113.6181 |
| 42 | 09-May-21 | 5 | M | -22.7435 | 113.6181 |
| 45 | 09-May-21 | 6 | M | -22.7556 | 113.6221 |
| 46 | 09-May-21 | 6 | M | -22.7644 | 113.6251 |
| 51 | 10-May-21 | 8 | M | -22.7670 | 113.6305 |
| 53 | 10-May-21 | 5 | F | -22.7616 | 113.6065 |
| 55 | 10-May-21 | 6 | U | -22.7716 | 113.6318 |
| 56 | 10-May-21 | 8 | M | -22.7654 | 113.6260 |
| 63 | 11-May-21 | 4.5 | F | -22.8040 | 113.6425 |
| 64 | 11-May-21 | 3.5 | F | -22.7998 | 113.6457 |
| 70 | 11-May-21 | 6.5 | M | -22.8049 | 113.6496 |
| 72 | 11-May-21 | 6 | M | -22.8120 | 113.6611 |
| 73 | 11-May-21 | 5.5 | M | -22.8202 | 113.6631 |

**Table S2.**
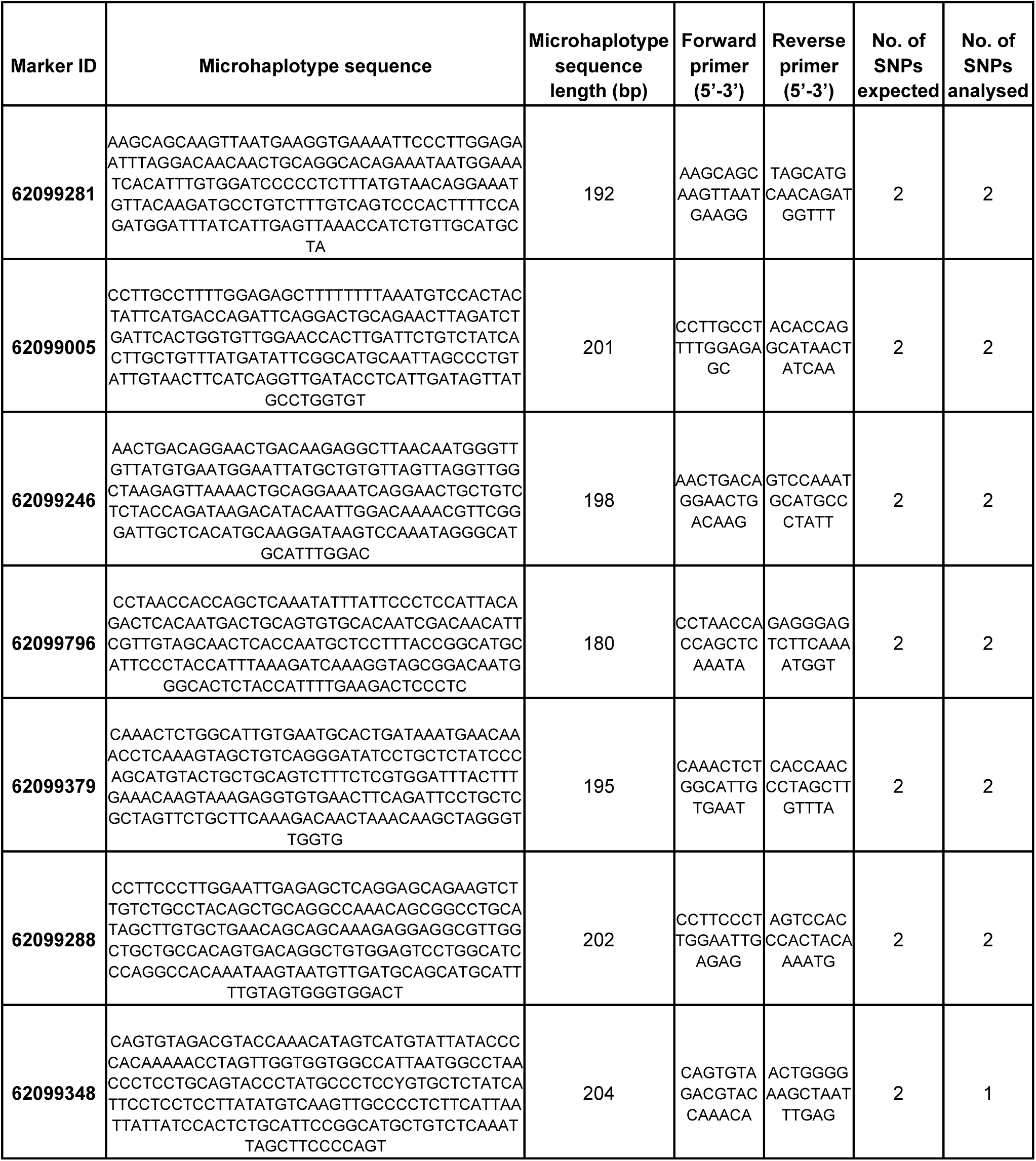
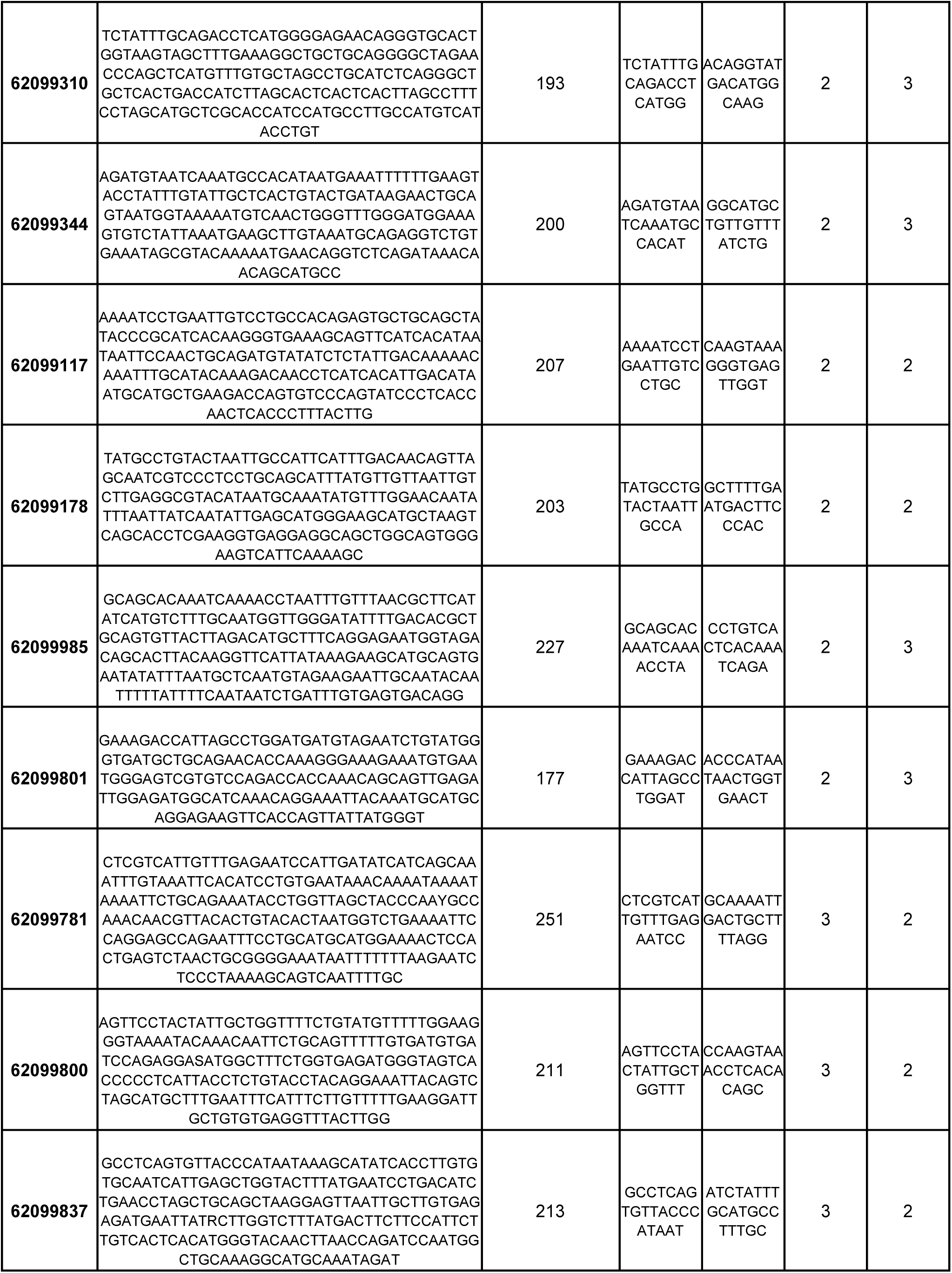
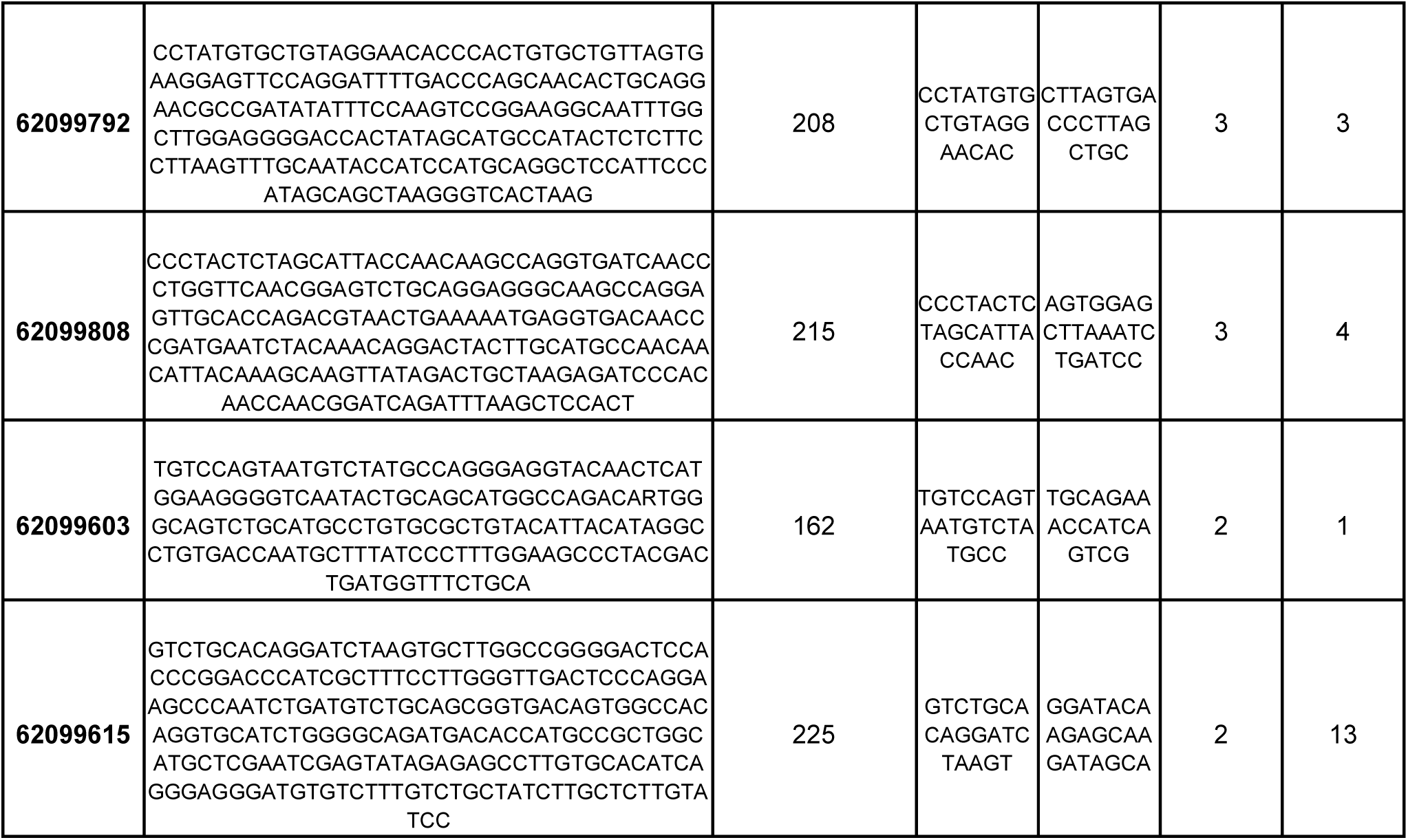
Metadata of 20 target microhaplotype sequences and primers used for analyses along with the number of SNPs expected according to DArT sequence data versus the actual number of SNPs obtained by analysing multiplex PCR sequence data.

**Figure S1.**
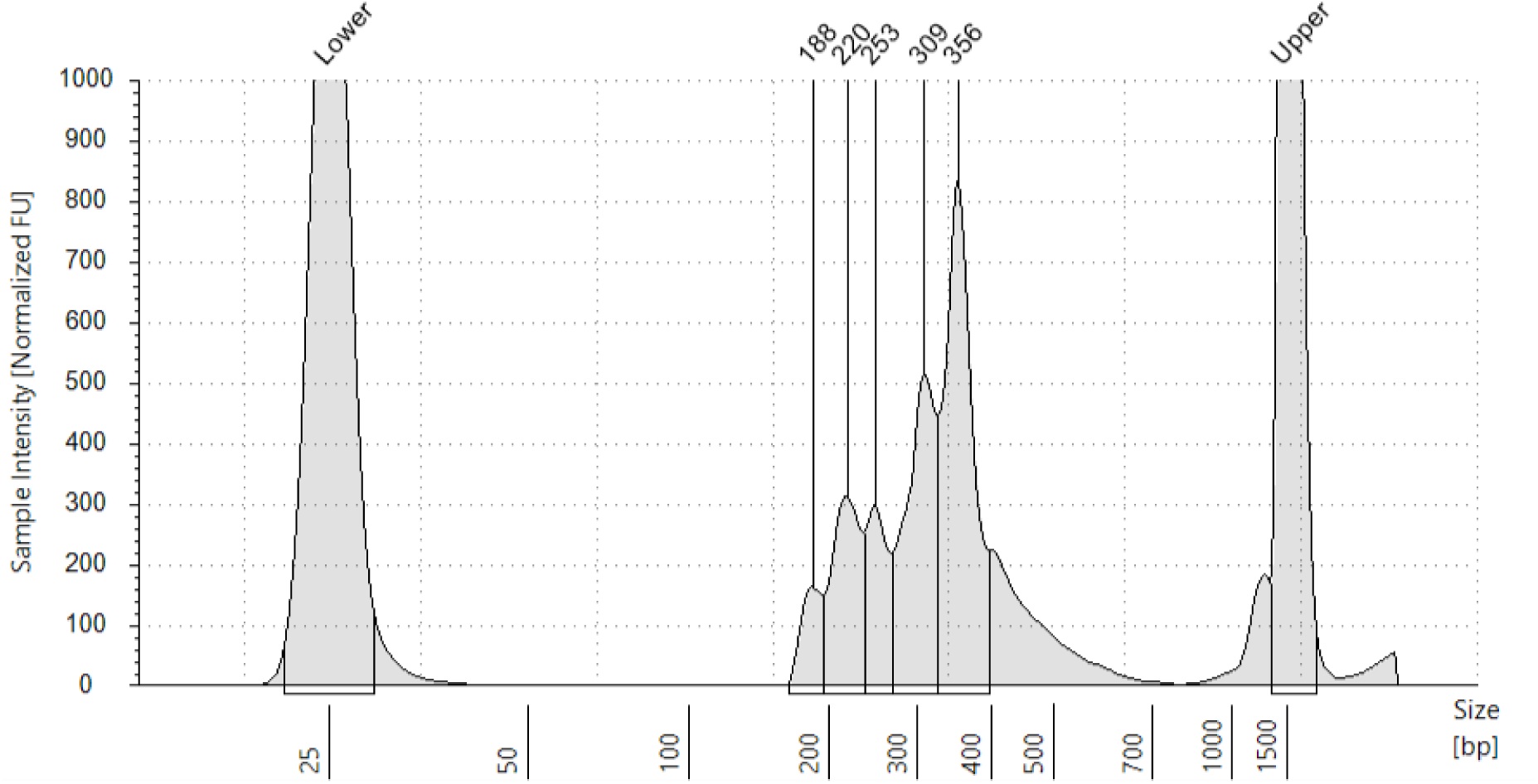
TapeStation DNA fragment analysis peak image showing size (base pairs) and concentration of fragments from a multiplexed, Indexed and size selected library pool of 96 whale shark tissue & eDNA samples ready to be sequenced.

**Figure S2.**
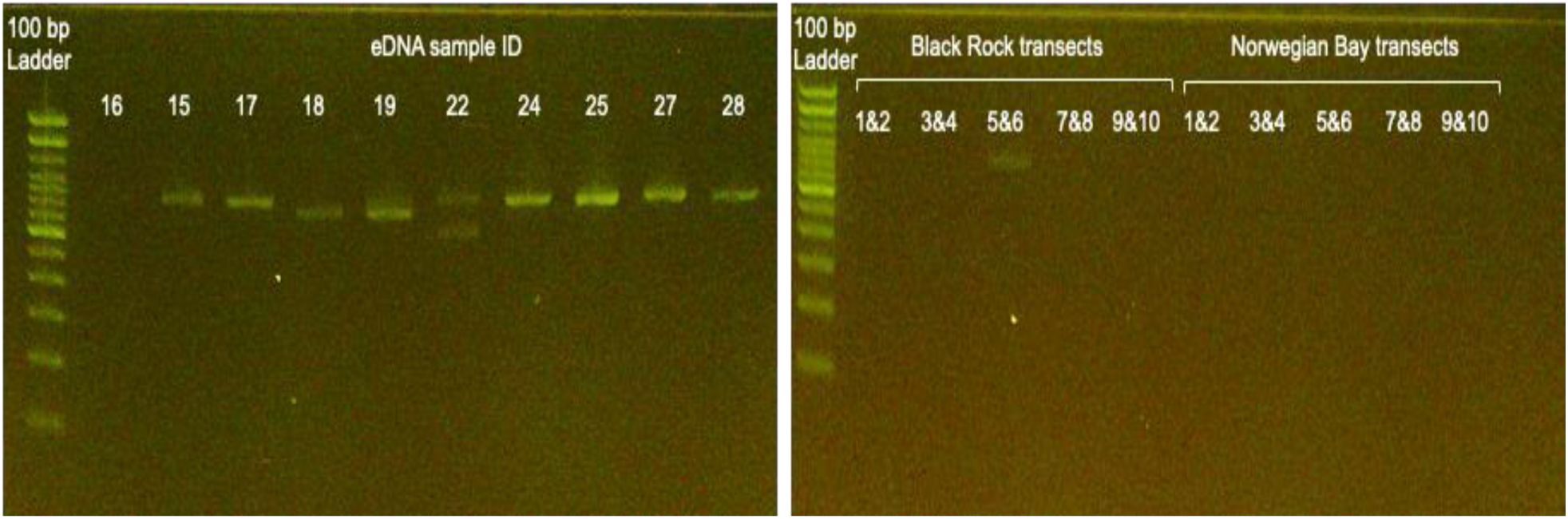
E-gel electrophoresis images of PCR amplicons (700 bp) of the WSCR whale shark mitochondrial control region used to screen eDNA samples for presence of whale shark mitochondrial DNA. The gel on the left shows eDNA samples collected behind individual sharks with all samples except sample 16 showing target amplification. The gel image on the right shows screening of transect eDNA samples, with only one sample from Black Rock transect 5&6 showing low amounts of WSCR amplification.

**Figure S3.**
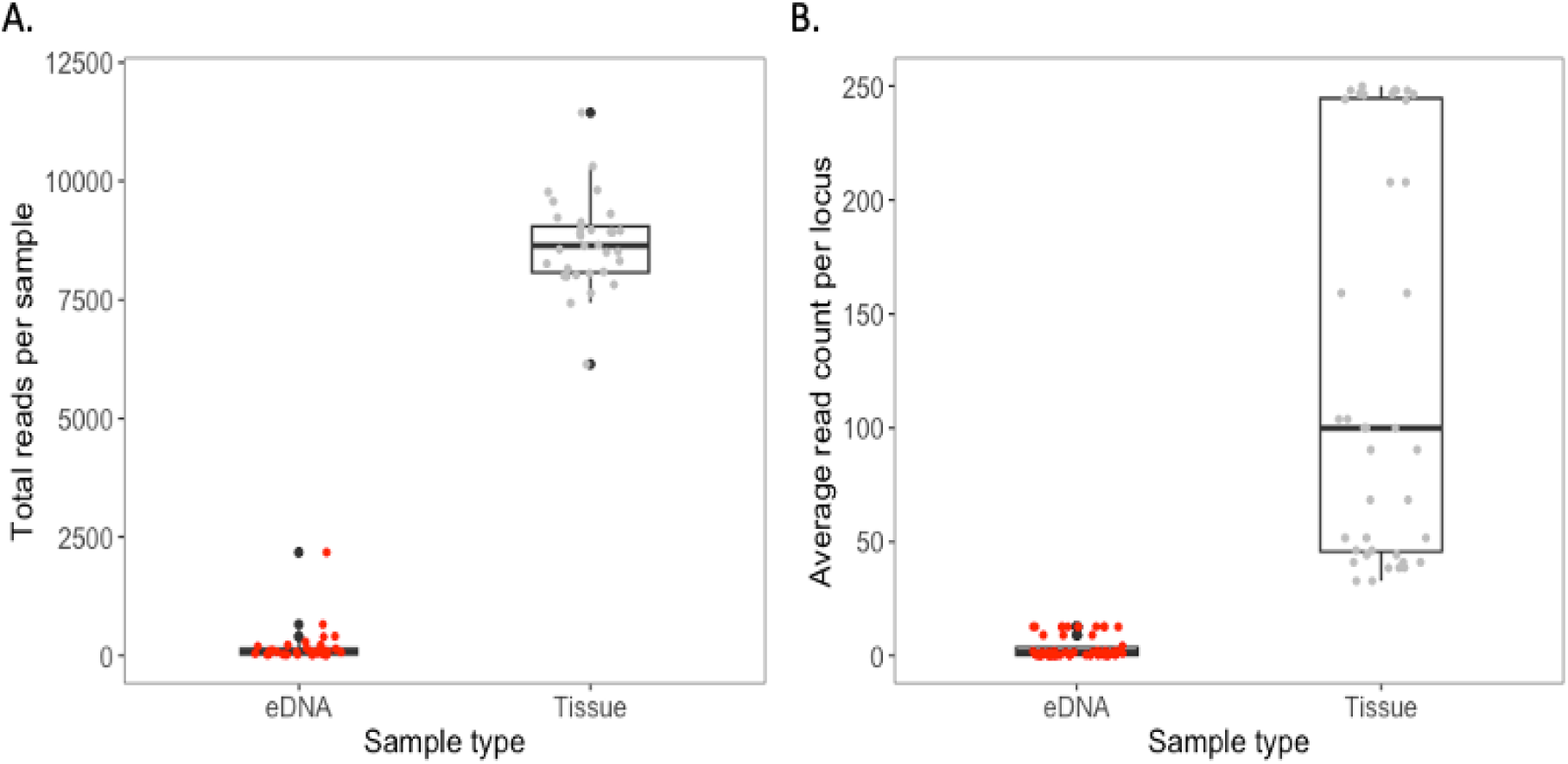
A. Total read counts obtained for each eDNA and tissue sample. B. Average read counts per locus across all eDNA and tissue samples.

**Figure S4.**
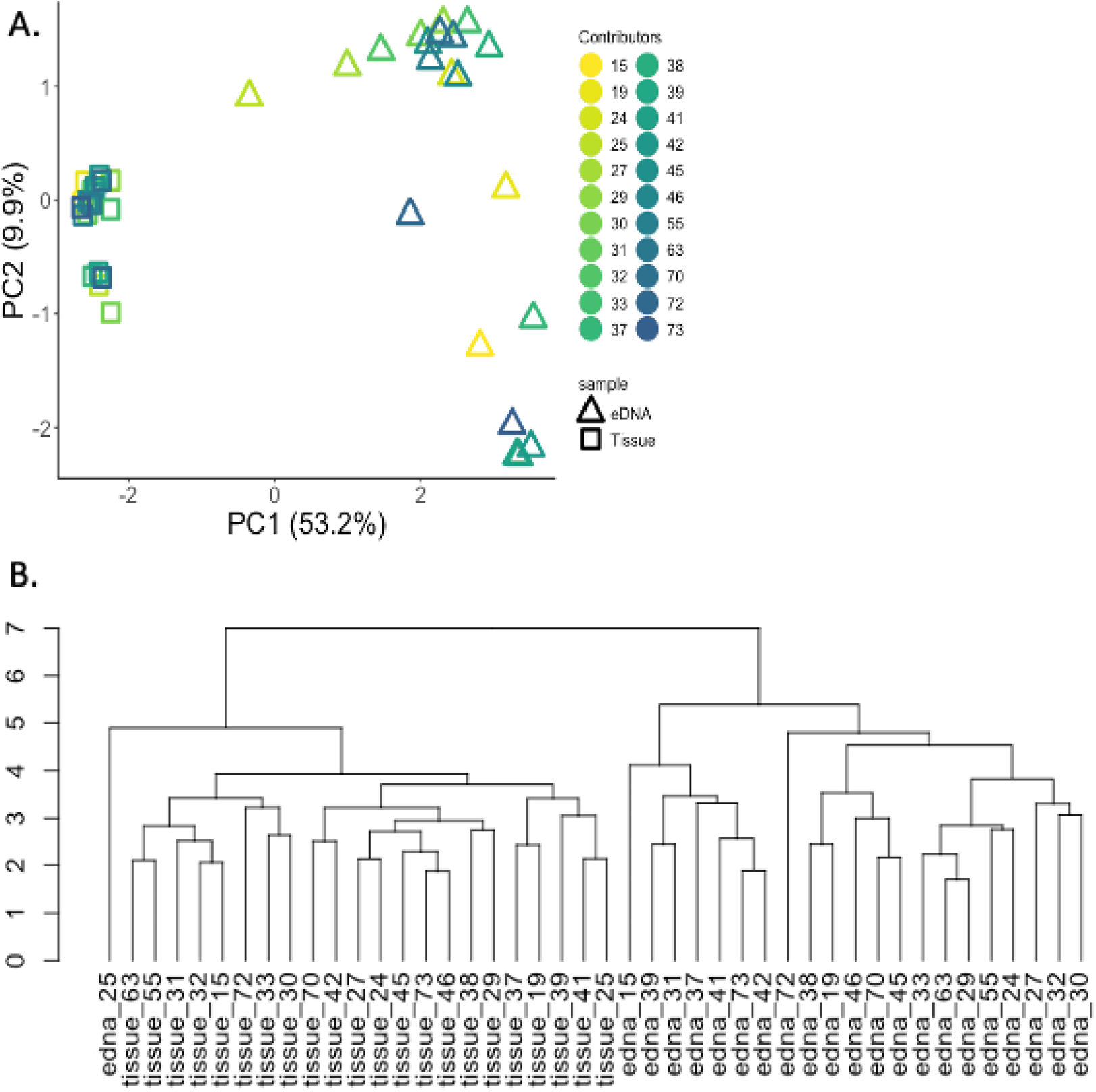
Assessing genetic similarity between eDNA and tissue samples for each individual whale shark by comparing allele frequencies, with all 112 alleles included. (A) Principal Component Analysis (PCA) plot illustrating individual eDNA and tissue samples, color-coded by sample ID. Solid black lines connect paired samples that belong to the same whale shark. (B) Dendrogram constructed using Euclidean distances to highlight clustering patterns between eDNA and tissue samples.

**Table S3.** Genetic diversity estimates for the whale shark population attending the Ningaloo Reef aggregation from 2016 to 2021, assessed using SNPs from a microhaplotype panel and genome-wide SNPs obtained via low-coverage Whole Genome Sequencing (lcWGS) (Meenakshisundaram et al., 2025b). For comparison, genetic diversity estimates from microhaplotype markers sequenced from environmental DNA (eDNA) samples collected in 2021 are also included alongside tissue-derived statistics.

|  | 2016 | 2017 | 2018 | 2019 | 2020 | 2021 | 2021-eDNA |
| --- | --- | --- | --- | --- | --- | --- | --- |
| Diversity estimates using microhaplotype panel: (number of SNP sites: 48) |  |  |  |  |  |  |  |
| No. of Individuals (N) | 10 | 10 | 10 | 10 | 10 | 22 | 22 |
| Observed heterozygosity (Ho) | 0.171 (0.18) | 0.202 (0.21) | 0.151 (0.19) | 0.206 (0.24) | 0.231 (0.22) | 0.194 (0.19) | 0.194 (0.16) |
| Unbiased expected heterozygosity (uHe) | 0.158 (0.17) | 0.187 (0.18) | 0.142 (0.17) | 0.177 (0.17) | 0.196 (0.17) | 0.182 (0.17) | 0.173 (0.13) |
| Allelic Richness (AR) | 1.154 (0.16) | 1.181 (0.17) | 1.139 (0.17) | 1.175 (0.17) | 1.191 (0.16) | 1.180 (0.17) | 1.171 (0.13) |
| Inbreeding coefficient (Fis) | -0.081 | -0.083 | -0.059 | -0.168 | -0.181 | -0.067 | -0.121 |
| Diversity estimates from lcWGS: (number of SNP sites: 7585) |  |  |  |  |  |  |  |
| No. of Individuals (N) | 10 | 10 | 10 | 16 | 10 | 8 | - |
| Observed heterozygosity (Ho) | 0.222 (.15) | 0.221 (.15) | 0.220 (.15) | 0.202 (.14) | 0.222 (.15) | 0.218 (.16) | - |
| Unbiased expected heterozygosity (uHe) | 0.196 (.12) | 0.196 (.12) | 0.194 (.12) | 0.178 (.11) | 0.196 (.12) | 0.193 (.13) | - |
| Allelic Richness (AR) | 1.827 (.31) | 1.834 (.3) | 1.831 (.3) | 1.77 (.27) | 1.834 (.3) | 1.829 (.38) | - |
| Inbreeding coefficient (Fis) | - 0.135 | - 0.132 | - 0.131 | - 0.136 | - 0.133 | - 0.128 | - |

## References

1. Acuña-Marrero, D., Jiménez, J., Smith, F., Doherty, P. F., Hearn, A., Green, J. R., Paredes-Jarrín, J., & Salinas-de-León, P. (2014). Whale Shark (Rhincodon typus) Seasonal Presence, Residence Time and Habitat Use at Darwin Island, Galapagos Marine Reserve. PLoS ONE, 9(12), e115946. 10.1371/journal.pone.0115946

2. Adamack, A. T., & Gruber, B. (2014). P OP G EN R EPORT: Simplifying basic population genetic analyses in R. Methods in Ecology and Evolution, 5(4), 384–387. 10.1111/2041-210X.12158

3. Adams, C. I. M., Knapp, M., Gemmell, N. J., Jeunen, G.-J., Bunce, M., Lamare, M. D., & Taylor, H. R. (2019). Beyond Biodiversity: Can Environmental DNA (eDNA) Cut It as a Population Genetics Tool? Genes, 10(3), 192. 10.3390/genes10030192

4. Adrian-Kalchhauser, I., & Burkhardt-Holm, P. (2016). An eDNA Assay to Monitor a Globally Invasive Fish Species from Flowing Freshwater. PLOS ONE, 11(1), e0147558. 10.1371/journal.pone.0147558

5. Alexander, J. B., Marnane, M. J., McDonald, J. I., Lukehurst, S. S., Elsdon, T. S., Simpson, T., Hinz, S., Bunce, M., & Harvey, E. S. (2023). Comparing environmental DNA collection methods for sampling community composition on marine infrastructure. *Estuarine*, Coastal and Shelf Science, 283, 108283. 10.1016/j.ecss.2023.108283

6. Ali Abd AlHameed, K. (2022). Spearmans correlation coefficient in statistical analysis. International Journal of Nonlinear Analysis and Applications, 13(1). 10.22075/ijnaa.2022.6079

7. Allendorf, F. W., Funk, W. C., Aitken, S. N., Byrne, M., & Luikart, G. (2022). Population Genomics. In F. W. Allendorf, W. C. Funk, S. N. Aitken, M. Byrne, G. Luikart, & A. Antunes, Conservation and the Genomics of Populations (3rd ed., pp. 66–92). Oxford University Press: Oxford. 10.1093/oso/9780198856566.003.0004

8. Andres, K. J., Lodge, D. M., Sethi, S. A., & Andrés, J. (2023). Detecting and analysing intraspecific genetic variation with EDNA: From population genetics to species abundance. Molecular Ecology, 32(15), 4118–4132. 10.1111/mec.17031

9. Andres, K. J., Sethi, S. A., Lodge, D. M., & Andrés, J. (2021). Nuclear eDNA estimates population allele frequencies and abundance in experimental mesocosms and field samples. Molecular Ecology, 30(3), 685–697. 10.1111/mec.15765

10. Arrowsmith, L., Sequeira, A., Pattiaratchi, C., & Meekan, M. (2021). Water temperature is a key driver of horizontal and vertical movements of an ocean giant, the whale shark Rhincodon typus. Marine Ecology Progress Series, 679, 101–114. 10.3354/meps13899

11. Baetscher, D. S., Beck, J., Anderson, E. C., Ruegg, K., Ramey, A. M., Hatch, S., Nevins, H., Fitzgerald, S. M., & Carlos Garza, J. (2022). Genetic assignment of fisheries bycatch reveals disproportionate mortality among Alaska Northern Fulmar breeding colonies. Evolutionary Applications, 15(3), 447–458. 10.1111/eva.13357

12. Baetscher, D. S., Clemento, A. J., Ng, T. C., Anderson, E. C., & Garza, J. C. (2018). Microhaplotypes provide increased power from short-read DNA sequences for relationship inference. Molecular Ecology Resources, 18(2), 296–305. 10.1111/1755-0998.12737

13. Beaumont, M. A. (2007). Conservation Genetics. In D. J. Balding, M. Bishop, & C. Cannings (Eds.), Handbook of Statistical Genetics (1st ed., pp. 1021–1066). Wiley. 10.1002/9780470061619.ch30

14. Berry, T. E., Saunders, B. J., Coghlan, M. L., Stat, M., Jarman, S., Richardson, A. J., Davies, C. H., Berry, O., Harvey, E. S., & Bunce, M. (2019). Marine environmental DNA biomonitoring reveals seasonal patterns in biodiversity and identifies ecosystem responses to anomalous climatic events. PLOS Genetics, 15(2), e1007943. 10.1371/journal.pgen.1007943

15. Boca, S. M., Huang, L., & Rosenberg, N. A. (2020). On the heterozygosity of an admixed population. Journal of Mathematical Biology, 81(6–7), 1217–1250. 10.1007/s00285-020-01531-9

16. Bradshaw, C. J. A., Mollet, H. F., & Meekan, M. G. (2007). Inferring population trends for the world’s largest fish from mark–recapture estimates of survival. Journal of Animal Ecology, 76(3), 480–489. 10.1111/j.1365-2656.2006.01201.x

17. Bylemans, J., Furlan, E. M., Gleeson, D. M., Hardy, C. M., & Duncan, R. P. (2018). Does Size Matter? An Experimental Evaluation of the Relative Abundance and Decay Rates of Aquatic Environmental DNA. Environmental Science & Technology, 52(11), 6408– 6416. 10.1021/acs.est.8b01071

18. Camacho, C., Coulouris, G., Avagyan, V., Ma, N., Papadopoulos, J., Bealer, K., & Madden, T. L. (2009). BLAST+: Architecture and applications. BMC Bioinformatics, 10(1), 421. 10.1186/1471-2105-10-421

19. Castro, A. L. F., Stewart, B. S., Wilson, S. G., Hueter, R. E., Meekan, M. G., Motta, P. J., Bowen, B. W., & Karl, S. A. (2007). Population genetic structure of Earth’s largest fish, the whale shark (*Rhincodon typus*). Molecular Ecology, 16(24), 5183–5192. 10.1111/j.1365-294X.2007.03597.x

20. Chen, P., Yin, C., Li, Z., Pu, Y., Yu, Y., Zhao, P., Chen, D., Liang, W., Zhang, L., & Chen, F. (2018). Evaluation of the Microhaplotypes panel for DNA mixture analyses. Forensic Science International: Genetics, 35, 149–155. 10.1016/j.fsigen.2018.05.003

21. Chen, S., Zhou, Y., Chen, Y., & Gu, J. (2018). fastp: An ultra-fast all-in-one FASTQ preprocessor. Bioinformatics, 34(17), i884–i890. 10.1093/bioinformatics/bty560

22. Collins, R. A., Wangensteen, O. S., O’Gorman, E. J., Mariani, S., Sims, D. W., & Genner, M. J. (2018). Persistence of environmental DNA in marine systems. Communications Biology, 1(1), 185. 10.1038/s42003-018-0192-6

23. Costello, M. J. (2022). Threats to Marine Species and Habitats, and How Banning Seabed Trawling Supports the Global Biodiversity Framework. In Imperiled: The Encyclopedia of Conservation (pp. 633–639). Elsevier. 10.1016/B978-0-12-821139-7.00246-4

24. Czech, L., Spence, J. P., & Expósito-Alonso, M. (2024). grenedalf: Population genetic statistics for the next generation of pool sequencing. Bioinformatics, 40(8), btae508. 10.1093/bioinformatics/btae508

25. D’Aloia, C. C., Andrés, J. A., Bogdanowicz, S. M., McCune, A. R., Harrison, R. G., & Buston, P. M. (2020). Unraveling hierarchical genetic structure in a marine metapopulation: A comparison of three high-throughput genotyping approaches. Molecular Ecology, 29(12), 2189–2203. 10.1111/mec.15405

26. Danecek, P., Auton, A., Abecasis, G., Albers, C. A., Banks, E., DePristo, M. A., Handsaker, R. E., Lunter, G., Marth, G. T., Sherry, S. T., McVean, G., Durbin, R., & 1000 Genomes Project Analysis Group. (2011). The variant call format and VCFtools. Bioinformatics, 27(15), 2156–2158. 10.1093/bioinformatics/btr330

27. Danecek, P., Bonfield, J. K., Liddle, J., Marshall, J., Ohan, V., Pollard, M. O., Whitwham, A., Keane, T., McCarthy, S. A., Davies, R. M., & Li, H. (2021). Twelve years of SAMtools and BCFtools. GigaScience, 10(2), giab008. 10.1093/gigascience/giab008

28. De La Parra Venegas, R., Hueter, R., González Cano, J., Tyminski, J., Gregorio Remolina, J., Maslanka, M., Ormos, A., Weigt, L., Carlson, B., & Dove, A. (2011). An Unprecedented Aggregation of Whale Sharks, Rhincodon typus, in Mexican Coastal Waters of the Caribbean Sea. PLoS ONE, 6(4), e18994. 10.1371/journal.pone.0018994

29. Dejean, T., Valentini, A., Duparc, A., Pellier-Cuit, S., Pompanon, F., Taberlet, P., & Miaud, C. (2011). Persistence of Environmental DNA in Freshwater Ecosystems. PLoS ONE, 6(8), e23398. 10.1371/journal.pone.0023398

30. Dekker, W. (2016). Management of the eel is slipping through our hands! Distribute control and orchestrate national protection. ICES Journal of Marine Science: Journal Du Conseil, 73(10), 2442–2452. 10.1093/icesjms/fsw094

31. Dugal, L., Thomas, L., Jensen, M. R., Sigsgaard, E. E., Simpson, T., Jarman, S., Thomsen, P. F., & Meekan, M. (2022a). Individual haplotyping of whale sharks from seawater environmental DNA. Molecular Ecology Resources, 22(1), 56–65. 10.1111/1755-0998.13451

32. Dugal, L., Thomas, L., Wilkinson, S. P., Richards, Z. T., Alexander, J. B., Adam, A. A. S., Kennington, W. J., Jarman, S., Ryan, N. M., Bunce, M., & Gilmour, J. P. (2022b). Coral monitoring in northwest Australia with environmental DNA metabarcoding using a curated reference database for optimized detection. Environmental DNA, 4(1), 63–76. 10.1002/edn3.199

33. Egeland, T., Dalen, I., & Mostad, P. F. (2003). Estimating the number of contributors to a DNA profile. International Journal of Legal Medicine, 117(5), 271–275. 10.1007/s00414-003-0382-7

34. Foote, A. D., Thomsen, P. F., Sveegaard, S., Wahlberg, M., Kielgast, J., Kyhn, L. A., Salling, A. B., Galatius, A., Orlando, L., & Gilbert, M. T. P. (2012). Investigating the Potential Use of Environmental DNA (eDNA) for Genetic Monitoring of Marine Mammals. PLoS ONE, 7(8), e41781. 10.1371/journal.pone.0041781

35. Fraija-Fernández, N., Bouquieaux, M., Rey, A., Mendibil, I., Cotano, U., Irigoien, X., Santos, M., & Rodríguez-Ezpeleta, N. (2020). Marine water environmental DNA metabarcoding provides a comprehensive fish diversity assessment and reveals spatial patterns in a large oceanic area. Ecology and Evolution, 10(14), 7560–7584. 10.1002/ece3.6482

36. G Carleton, Ray., Jerry, McCormick-Ray., Robert, L., & Smith. (2013). Marine Conservation: Science, Policy, and Management.

37. Gautier, M., Foucaud, J., Gharbi, K., Cézard, T., Galan, M., Loiseau, A., Thomson, M., Pudlo, P., Kerdelhué, C., & Estoup, A. (2013). Estimation of population allele frequencies from next-generation sequencing data: Pool-versus individual-based genotyping. Molecular Ecology, 22(14), 3766–3779. 10.1111/mec.12360

38. Gautier, M., Vitalis, R., Flori, L., & Estoup, A. (2022). *F* –Statistics estimation and admixture graph construction with Pool-Seq or allele count data using the R package *poolfstat*. Molecular Ecology Resources, 22(4), 1394–1416. 10.1111/1755-0998.13557

39. Germanov, E. S., Marshall, A. D., Bejder, L., Fossi, M. C., & Loneragan, N. R. (2018). Microplastics: No Small Problem for Filter-Feeding Megafauna. Trends in Ecology & Evolution, 33(4), 227–232. 10.1016/j.tree.2018.01.005

40. Gero, S., Milligan, M., Rinaldi, C., Francis, P., Gordon, J., Carlson, C., Steffen, A., Tyack, P., Evans, P., & Whitehead, H. (2014). Behavior and social structure of the sperm whales of Dominica, West Indies. Marine Mammal Science, 30(3), 905–922. 10.1111/mms.12086

41. Goldberg, C. S., Turner, C. R., Deiner, K., Klymus, K. E., Thomsen, P. F., Murphy, M. A., Spear, S. F., McKee, A., Oyler-McCance, S. J., Cornman, R. S., Laramie, M. B., Mahon, A. R., Lance, R. F., Pilliod, D. S., Strickler, K. M., Waits, L. P., Fremier, A. K., Takahara, T., Herder, J. E., & Taberlet, P. (2016). Critical considerations for the application of environmental DNA methods to detect aquatic species. Methods in Ecology and Evolution, 7(11), 1299–1307. 10.1111/2041-210X.12595

42. Gruber, B., Unmack, P. J., Berry, O. F., & Georges, A. (2018). DARTR: An R package to facilitate analysis of SNP data generated from reduced representation genome sequencing. Molecular Ecology Resources, 18(3), 691–699. 10.1111/1755-0998.12745

43. Hardenstine, R. S., He, S., Cochran, J. E. M., Braun, C. D., Cagua, E. F., Pierce, S. J., Prebble, C. E. M., Rohner, C. A., Saenz-Agudelo, P., Sinclair-Taylor, T. H., Skomal, G. B., Thorrold, S. R., Watts, A. M., Zakroff, C. J., & Berumen, M. L. (2022). Pieces in a global puzzle: Population genetics at two whale shark aggregations in the western Indian Ocean. Ecology and Evolution, 12(1), e8492. 10.1002/ece3.8492

44. Hearn, A. R., Green, J. R., Peñaherrera-Palma, C. R., Reynolds, S., Rohner, C. A., Román, M., & Sequeira, A. M. M. (2021). Whale Shark Movements and Migrations. In A. D. M. Dove & S. J. Pierce, Whale Sharks (1st ed., pp. 105–127). CRC Press. 10.1201/b22502-6

45. Hopken, M. W., Piaggio, A. J., Abdo, Z., Chipman, R. B., Mankowski, C. P., Nelson, K. M., Hilton, M. S., Thurber, C., Tsuchiya, M. T. N., Maldonado, J. E., & Gilbert, A. T. (2023). Are rabid raccoons (*Procyon lotor*) ready for the rapture? Determining the geographic origin of rabies virus-infected raccoons using RADCAPTURE and microhaplotypes. Evolutionary Applications, 16(12), 1937–1955. 10.1111/eva.13613

46. Hughes-Stamm, S. R. J., Ashton K., & Van Daal, A. (2011). Assessment of DNA degradation and the genotyping success of highly degraded samples. International Journal of Legal Medicine, 125(3), 341–348. 10.1007/s00414-010-0455-3

47. Huveneers, C., Meekan, M. G., Apps, K., Ferreira, L. C., Pannell, D., & Vianna, G. M. S. (2017). The economic value of shark-diving tourism in Australia. Reviews in Fish Biology and Fisheries, 27(3), 665–680. 10.1007/s11160-017-9486-x

48. IUCN. (2016). Rhincodon typus: Pierce, S.J. & Norman, B.: The IUCN Red List of Threatened Species 2016: *e.T19488A2365291* [Dataset]. 10.2305/IUCN.UK.2016-1.RLTS.T19488A2365291.en

49. James, M. C., Myers, R. A., & Ottensmeyer, C. A. (2005). Behaviour of leatherback sea turtles, Dermochelys coriacea, during the migratory cycle. Proceedings of the Royal Society B: Biological Sciences, 272(1572), 1547–1555. 10.1098/rspb.2005.3110

50. Jo, T., Takao, K., & Minamoto, T. (2022). Linking the state of environmental DNA to its application for biomonitoring and stock assessment: Targeting mitochondrial/nuclear genes, and different DNA fragment lengths and particle sizes. Environmental DNA, 4(2), 271–283. 10.1002/edn3.253

51. Johnson, M., Zaretskaya, I., Raytselis, Y., Merezhuk, Y., McGinnis, S., & Madden, T. L. (2008). NCBI BLAST: A better web interface. Nucleic Acids Research, 36(Web Server), W5– W9. 10.1093/nar/gkn201

52. Kechin, A., Borobova, V., Boyarskikh, U., Khrapov, E., Subbotin, S., & Filipenko, M. (2020). NGS-PrimerPlex: High-throughput primer design for multiplex polymerase chain reactions. PLOS Computational Biology, 16(12), e1008468. 10.1371/journal.pcbi.1008468

53. Ketchum, J. T., Galván-Magaña, F., & Klimley, A. P. (2013). Segregation and foraging ecology of whale sharks, Rhincodon typus, in the southwestern Gulf of California. Environmental Biology of Fishes, 96(6), 779–795. 10.1007/s10641-012-0071-9

54. Kidd, K. K., Pakstis, A. J., Speed, W. C., Lagacé, R., Chang, J., Wootton, S., Haigh, E., & Kidd, J. R. (2014). Current sequencing technology makes microhaplotypes a powerful new type of genetic marker for forensics. Forensic Science International: Genetics, 12, 215–224. 10.1016/j.fsigen.2014.06.014

55. Kidd, K. K., Pakstis, A. J., Speed, W. C., Lagace, R., Chang, J., Wootton, S., & Ihuegbu, N. (2013). Microhaplotype loci are a powerful new type of forensic marker. Forensic Science International: Genetics Supplement Series, 4(1), e123–e124. 10.1016/j.fsigss.2013.10.063

56. Kidd, K. K., & Speed, W. C. (2015). Criteria for selecting microhaplotypes: Mixture detection and deconvolution. Investigative Genetics, 6(1), 1. 10.1186/s13323-014-0018-3

57. Kumar, G., Eble, J. E., & Gaither, M. R. (2020). A practical guide to sample preservation and pre-PCR processing of aquatic environmental DNA. Molecular Ecology Resources, 20(1), 29–39. 10.1111/1755-0998.13107

58. Lester, E., Meekan, M., Barnes, P., Raudino, H., Rob, D., Waples, K., & Speed, C. (2020). Multi-year patterns in scarring, survival and residency of whale sharks in Ningaloo Marine Park, Western Australia. Marine Ecology Progress Series, 634, 115–125. 10.3354/meps13173

59. Lewison, R. L., Johnson, A. F., & Verutes, G. M. (2018). Embracing Complexity and Complexity-Awareness in Marine Megafauna Conservation and Research. Frontiers in Marine Science, 5, 207. 10.3389/fmars.2018.00207

60. Li, H. (2013). *Aligning sequence reads, clone sequences and assembly contigs with BWA-MEM* (arXiv:1303.3997). arXiv. 10.48550/arXiv.1303.3997

61. Li, H., Handsaker, B., Wysoker, A., Fennell, T., Ruan, J., Homer, N., Marth, G., Abecasis, G., Durbin, R., & 1000 Genome Project Data Processing Subgroup. (2009). The Sequence Alignment/Map format and SAMtools. Bioinformatics, 25(16), 2078–2079. 10.1093/bioinformatics/btp352

62. Lippé, C., Dumont, P., & Bernatchez, L. (2006). High genetic diversity and no inbreeding in the endangered copper redhorse, *Moxostoma hubbsi* (Catostomidae, Pisces): The positive sides of a long generation time. Molecular Ecology, 15(7), 1769–1780. 10.1111/j.1365-294X.2006.02902.x

63. Majaneva, M., Diserud, O. H., Eagle, S. H. C., Boström, E., Hajibabaei, M., & Ekrem, T. (2018). Environmental DNA filtration techniques affect recovered biodiversity. Scientific Reports, 8(1), 4682. 10.1038/s41598-018-23052-8

64. Marshall, C., & Parson, W. (2023). Mitochondrial DNA. In Encyclopedia of Forensic Sciences, Third Edition (pp. 592–601). Elsevier. 10.1016/B978-0-12-823677-2.00138-0

65. McClain, C. R., Balk, M. A., Benfield, M. C., Branch, T. A., Chen, C., Cosgrove, J., Dove, A. D. M., Gaskins, L., Helm, R. R., Hochberg, F. G., Lee, F. B., Marshall, A., McMurray, S. E., Schanche, C., Stone, S. N., & Thaler, A. D. (2015). Sizing ocean giants: Patterns of intraspecific size variation in marine megafauna. PeerJ, 3, e715. 10.7717/peerj.715

66. Meekan, M., Austin, C. M., Tan, M. H., Wei, N.-W. V., Miller, A., Pierce, S. J., Rowat, D., Stevens, G., Davies, T. K., Ponzo, A., & Gan, H. M. (2017). iDNA at Sea: Recovery of Whale Shark (Rhincodon typus) Mitochondrial DNA Sequences from the Whale Shark Copepod (Pandarus rhincodonicus) Confirms Global Population Structure. Frontiers in Marine Science, 4, 420. 10.3389/fmars.2017.00420

67. Meekan, M., Bradshaw, C., Press, M., McLean, C., Richards, A., Quasnichka, S., & Taylor, J. (2006). Population size and structure of whale sharks Rhincodon typus at Ningaloo Reef, Western Australia. Marine Ecology Progress Series, 319, 275–285. 10.3354/meps319275

68. Meekan, M. G., Taylor, B. M., Lester, E., Ferreira, L. C., Sequeira, A. M. M., Dove, A. D. M., Birt, M. J., Aspinall, A., Brooks, K., & Thums, M. (2020). Asymptotic Growth of Whale Sharks Suggests Sex-Specific Life-History Strategies. Frontiers in Marine Science, 7, 575683. 10.3389/fmars.2020.575683

69. Meenakshisundaram, A., Jarman, S., Power, H., Kennington, W. J., & Thomas, L. (2025). Population genetics in aquatic environments using environmental DNA microhaplotypes: A case study of zebrafish in controlled aquaria. (Preprint in www.biorxiv.org)

70. Meenakshisundaram, A., Meekan, M., Kaur, P., Pearce, J., Kennington, W. J., & Thomas, L. (under prep.). A genomic analysis of the whale shark, Rhincodon typus, through contemporary and evolutionary time.

71. Meenakshisundaram, A., Thomas, L., Kennington, W., Thums, M., Lester, E., & Meekan, M. (2021). Genetic markers validate photo-identification and uniqueness of spot patterns in whale sharks. Marine Ecology Progress Series, 668, 177–183. 10.3354/meps13729

72. Miya, M., Minamoto, T., Yamanaka, H., Oka, S., Sato, K., Yamamoto, S., Sado, T., & Doi, H. (2016). Use of a Filter Cartridge for Filtration of Water Samples and Extraction of Environmental DNA. Journal of Visualized Experiments, 117, 54741. 10.3791/54741

73. Moezi, P., Kargar, M., Doosti, A., & Khoshneviszadeh, M. (2019). Multiplex touchdown PCR assay to enhance specificity and sensitivity for concurrent detection of four foodborne pathogens in raw milk. Journal of Applied Microbiology, 127(1), 262–273. 10.1111/jam.14285

74. Morin, P. A., Forester, B. R., Forney, K. A., Crossman, C. A., Hancock-Hanser, B. L., Robertson, K. M., Barrett-Lennard, L. G., Baird, R. W., Calambokidis, J., Gearin, P., Hanson, M. B., Schumacher, C., Harkins, T., Fontaine, M. C., Taylor, B. L., & Parsons, K. M. (2021). Population structure in a continuously distributed coastal marine species, the harbor porpoise, based on microhaplotypes derived from poor-quality samples. Molecular Ecology, 30(6), 1457–1476. 10.1111/mec.15827

75. Moushomi, R., Wilgar, G., Carvalho, G., Creer, S., & Seymour, M. (2019). Environmental DNA size sorting and degradation experiment indicates the state of Daphnia magna mitochondrial and nuclear eDNA is subcellular. Scientific Reports, 9(1), 12500. 10.1038/s41598-019-48984-7

76. Murakami, H., Yoon, S., Kasai, A., Minamoto, T., Yamamoto, S., Sakata, M. K., Horiuchi, T., Sawada, H., Kondoh, M., Yamashita, Y., & Masuda, R. (2019). Dispersion and degradation of environmental DNA from caged fish in a marine environment. Fisheries Science, 85(2), 327–337. 10.1007/s12562-018-1282-6

77. Nevers, M. B., Byappanahalli, M. N., Morris, C. C., Shively, D., Przybyla-Kelly, K., Spoljaric, A. M., Dickey, J., & Roseman, E. F. (2018). Environmental DNA (eDNA): A tool for quantifying the abundant but elusive round goby (Neogobius melanostomus). PLOS ONE, 13(1), e0191720. 10.1371/journal.pone.0191720

78. O’Hara, C., Frazier, M., Valle, M., Butt, N., Kaschner, K., Klein, C., & Halpern, B. (2023). *Cumulative human impacts on global marine fauna highlight risk to fragile functional diversity of marine ecosystems* [Preprint]. Preprints. 10.22541/au.168328097.74947503/v1

79. Oldoni, F., Bader, D., Fantinato, C., Wootton, S. C., Lagacé, R., Kidd, K. K., & Podini, D. (2020). A sequence-based 74plex microhaplotype assay for analysis of forensic DNA mixtures. Forensic Science International: Genetics, 49, 102367. 10.1016/j.fsigen.2020.102367

80. Oldoni, F., Hart, R., Long, K., Maddela, K., Cisana, S., Schanfield, M., Wootton, S., Chang, J., Lagace, R., Hasegawa, R., Kidd, K., & Podini, D. (2017). Microhaplotypes for ancestry prediction. Forensic Science International: Genetics Supplement Series, 6, e513–e515. 10.1016/j.fsigss.2017.09.209

81. Oldoni, F., Kidd, K. K., & Podini, D. (2019). Microhaplotypes in forensic genetics. Forensic Science International: Genetics, 38, 54–69. 10.1016/j.fsigen.2018.09.009

82. Oleksiak, M. F., & Rajora, O. P. (2019). Marine Population Genomics: Challenges and Opportunities. In M. F. Oleksiak & O. P. Rajora (Eds.), Population Genomics: Marine Organisms (pp. 3–35). Springer International Publishing. 10.1007/13836_2019_70

83. Parsons, K. M., Everett, M., Dahlheim, M., & Park, L. (2018). Water, water everywhere: Environmental DNA can unlock population structure in elusive marine species. Royal Society Open Science, 5(8), 180537. 10.1098/rsos.180537

84. Pimiento, C., Leprieur, F., Silvestro, D., Lefcheck, J. S., Albouy, C., Rasher, D. B., Davis, M., Svenning, J.-C., & Griffin, J. N. (2020). Functional diversity of marine megafauna in the Anthropocene. Science Advances, 6(16), eaay7650. 10.1126/sciadv.aay7650

85. Quinlan, A. R. (2014). BEDTools: The Swiss-Army Tool for Genome Feature Analysis. Current Protocols in Bioinformatics, 47(1). 10.1002/0471250953.bi1112s47

86. R Core Team. (2022). computing. R Foundation for Statistical Computing, Vienna, Austria. https://www.R-project.org/.

87. Reynolds, S. D., Norman, B. M., Beger, M., Franklin, C. E., & Dwyer, R. G. (2017). Movement, distribution and marine reserve use by an endangered migratory giant. Diversity and Distributions, 23(11), 1268–1279. 10.1111/ddi.12618

88. Riley, M., Hale, M., Harman, A., & Rees, R. (2010). Analysis of whale shark Rhincodon typus aggregations near South Ari Atoll, Maldives Archipelago. Aquatic Biology, 8, 145–150. 10.3354/ab00215

89. Rognes, T., Flouri, T., Nichols, B., Quince, C., & Mahé, F. (2016). VSEARCH: A versatile open source tool for metagenomics. PeerJ, 4, e2584. 10.7717/peerj.2584

90. Rohner, C. A., Norman, B. M., Reynolds, S., Araujo, G., Holmberg, J., & Pierce, S. J. (2021). Population Ecology of Whale Sharks. In A. D. M. Dove & S. J. Pierce, Whale Sharks (1st ed., pp. 129–152). CRC Press. 10.1201/b22502-7

91. Rosenbaum, H. C., Pomilla, C., Mendez, M., Leslie, M. S., Best, P. B., Findlay, K. P., Minton, G., Ersts, P. J., Collins, T., Engel, M. H., Bonatto, S. L., Kotze, D. P. G. H., Meÿer, M., Barendse, J., Thornton, M., Razafindrakoto, Y., Ngouessono, S., Vely, M., & Kiszka, J. (2009). Population Structure of Humpback Whales from Their Breeding Grounds in the South Atlantic and Indian Oceans. PLoS ONE, 4(10), e7318. 10.1371/journal.pone.0007318

92. Rowat, D., Womersley, F., Norman, B. M., & Pierce, S. J. (2021). Global Threats to Whale Sharks. In A. D. M. Dove & S. J. Pierce, Whale Sharks (1st ed., pp. 239–265). CRC Press. 10.1201/b22502-11

93. Rus Hoelzel, A., Shivji, M. S., Magnussen, J., & Francis, M. P. (2006). Low worldwide genetic diversity in the basking shark (*Cetorhinus maximus*). Biology Letters, 2(4), 639–642. 10.1098/rsbl.2006.0513

94. Schmidt, J. V., Schmidt, C. L., Ozer, F., Ernst, R. E., Feldheim, K. A., Ashley, M. V., & Levine, M. (2009). Low Genetic Differentiation across Three Major Ocean Populations of the Whale Shark, Rhincodon typus. PLoS ONE, 4(4), e4988. 10.1371/journal.pone.0004988

95. Sea K., G., M., R., T., U., N. V., S. K., P., J., Keer, N. R., Yadav, R., & Yadav, A. K. (2022). Evaluating and ranking the Vulnerability of the marine ecosystem to multiple threats. The Indian Journal of Animal Sciences, 92(5), 654–658. 10.56093/ijans.v92i5.113397

96. Sethi, S. A., Larson, W., Turnquist, K., & Isermann, D. (2019). Estimating the number of contributors to DNA mixtures provides a novel tool for ecology. Methods in Ecology and Evolution, 10(1), 109–119. 10.1111/2041-210X.13079

97. Shirazi, S., Meyer, R., & Shapiro, B. (2021). Revisiting the effect of PCR replication and sequencing depth on biodiversity metrics in environmental DNA metabarcoding. 10.22541/au.159309876.62184178/v2

98. Shokolenko, I., Venediktova, N., Bochkareva, A., Wilson, G. L., & Alexeyev, M. F. (2009). Oxidative stress induces degradation of mitochondrial DNA. Nucleic Acids Research, 37(8), 2539–2548. 10.1093/nar/gkp100

99. Sigsgaard, E. E., Jensen, M. R., Winkelmann, I. E., Møller, P. R., Hansen, M. M., & Thomsen, P. F. (2020a). Population-level inferences from environmental DNA—Current status and future perspectives. Evolutionary Applications, 13(2), 245–262. 10.1111/eva.12882

100. Sigsgaard, E. E., Nielsen, I. B., Bach, S. S., Lorenzen, E. D., Robinson, D. P., Knudsen, S. W., Pedersen, M. W., Jaidah, M. A., Orlando, L., Willerslev, E., Møller, P. R., & Thomsen, P. F. (2016). Population characteristics of a large whale shark aggregation inferred from seawater environmental DNA. Nature Ecology & Evolution, 1(1), 0004. 10.1038/s41559-016-0004

101. Sigsgaard, E. E., Torquato, F., Frøslev, T. G., Moore, A. B. M., Sørensen, J. M., Range, P., Ben-Hamadou, R., Bach, S. S., Møller, P. R., & Thomsen, P. F. (2020b). Using vertebrate environmental DNA from seawater in biomonitoring of marine habitats. Conservation Biology, 34(3), 697–710. 10.1111/cobi.13437

102. Staadig, A., Hedman, J., & Tillmar, A. (2023). Applying Unique Molecular Indices with an Extensive All-in-One Forensic SNP Panel for Improved Genotype Accuracy and Sensitivity. Genes, 14(4), 818. 10.3390/genes14040818

103. Suarez-Bregua, P., Álvarez-González, M., Parsons, K. M., Rotllant, J., Pierce, G. J., & Saavedra, C. (2022). Environmental DNA (eDNA) for monitoring marine mammals: Challenges and opportunities. Frontiers in Marine Science, 9, 987774. 10.3389/fmars.2022.987774

104. Taberlet, P., Bonin, A., Zinger, L., & Coissac, E. (2018). Environmental DNA: For Biodiversity Research and Monitoring (1st ed.). Oxford University PressOxford. 10.1093/oso/9780198767220.001.0001

105. Taberlet, P., & Luikart, G. (1999). Non-invasive genetic sampling and individual identification. Biological Journal of the Linnean Society, 68(1–2), 41–55. 10.1111/j.1095-8312.1999.tb01157.x

106. Takahashi, M., Saccò, M., Kestel, J. H., Nester, G., Campbell, M. A., Van Der Heyde, M., Heydenrych, M. J., Juszkiewicz, D. J., Nevill, P., Dawkins, K. L., Bessey, C., Fernandes, K., Miller, H., Power, M., Mousavi-Derazmahalleh, M., Newton, J. P., White, N. E., Richards, Z. T., & Allentoft, M. E. (2023). Aquatic environmental DNA: A review of the macro-organismal biomonitoring revolution. Science of The Total Environment, 873, 162322. 10.1016/j.scitotenv.2023.162322

107. Thomsen, P. F., Kielgast, J., Iversen, L. L., Møller, P. R., Rasmussen, M., & Willerslev, E. (2012). Detection of a Diverse Marine Fish Fauna Using Environmental DNA from Seawater Samples. PLoS ONE, 7(8), e41732. 10.1371/journal.pone.0041732

108. Tsuji, S., Ushio, M., Sakurai, S., Minamoto, T., & Yamanaka, H. (2017). Water temperature-dependent degradation of environmental DNA and its relation to bacterial abundance. PLOS ONE, 12(4), e0176608. 10.1371/journal.pone.0176608

109. Van Tienhoven, A. M., Den Hartog, J. E., Reijns, R. A., & Peddemors, V. M. (2007). A computer-aided program for pattern-matching of natural marks on the spotted raggedtooth shark *Carcharias taurus*. Journal of Applied Ecology, 44(2), 273–280. 10.1111/j.1365-2664.2006.01273.x

110. Vignaud, T. M., Maynard, J. A., Leblois, R., Meekan, M. G., Vázquez-Juárez, R., Ramírez-Macías, D., Pierce, S. J., Rowat, D., Berumen, M. L., Beeravolu, C., Baksay, S., & Planes, S. (2014). Genetic structure of populations of whale sharks among ocean basins and evidence for their historic rise and recent decline. Molecular Ecology, 23(10), 2590–2601. 10.1111/mec.12754

111. Welsh, A., Hill, T., Quinlan, H., Robinson, C., & May, B. (2008). Genetic Assessment of Lake Sturgeon Population Structure in the Laurentian Great Lakes. North American Journal of Fisheries Management, 28(2), 572–591. 10.1577/M06-184.1

112. Womersley, F. C., Humphries, N. E., Queiroz, N., Vedor, M., Da Costa, I., Furtado, M., Tyminski, J. P., Abrantes, K., Araujo, G., Bach, S. S., Barnett, A., Berumen, M. L., Bessudo Lion, S., Braun, C. D., Clingham, E., Cochran, J. E. M., De La Parra, R., Diamant, S., Dove, A. D. M., … Sims, D. W. (2022). Global collision-risk hotspots of marine traffic and the world’s largest fish, the whale shark. Proceedings of the National Academy of Sciences, 119(20), e2117440119. 10.1073/pnas.2117440119

113. Wong, M. K.-S., Nakao, M., & Hyodo, S. (2020). Field application of an improved protocol for environmental DNA extraction, purification, and measurement using Sterivex filter. Scientific Reports, 10(1), 21531. 10.1038/s41598-020-77304-7

114. Ye, J., Coulouris, G., Zaretskaya, I., Cutcutache, I., Rozen, S., & Madden, T. L. (2012). Primer-BLAST: A tool to design target-specific primers for polymerase chain reaction. BMC Bioinformatics, 13(1), 134. 10.1186/1471-2105-13-134

115. Yoccoz, N. G. (2012). The future of environmental DNA in ecology. Molecular Ecology, 21(8), 2031–2038. 10.1111/j.1365-294X.2012.05505.x

116. Zheng, X., Levine, D., Shen, J., Gogarten, S. M., Laurie, C., & Weir, B. S. (2012). A high-performance computing toolset for relatedness and principal component analysis of SNP data. Bioinformatics, 28(24), 3326–3328. 10.1093/bioinformatics/bts606

